# Microbially-competent skin organotypics reveal dysregulated AhR signalling in microbial dysbiosis

**DOI:** 10.1101/2025.07.01.662505

**Authors:** Esmé P. McPolin-Hall, Abish S. Stephen, Claire Pardieu, Garrick Georgeu, Robert P. Allaker, Ranjit K. Bhogal, Jenny E. Pople, Michael P. Philpott, Rosalind F. Hannen

## Abstract

Microbial dysbiosis is increasingly linked to skin disorders, yet the mechanisms driving these pathological processes remain poorly understood. This study developed microbially competent 3D skin organotypics (OTs) representing healthy (5M) and dandruff (5MP) microbiomes, to investigate host-microbial interactions. OTs colonised with 5M microbiota exhibited healthy skin morphology, while 5MP colonised OTs displayed dandruff-like phenotypes, including altered epidermal morphology, reduced barrier protein expression, and dysregulated corneodesmosome hydrolysis. Transcriptomic and protein analysis revealed attenuation of the aryl hydrocarbon receptor (AhR) signalling pathway in 5MP colonised OTs, a finding subsequently validated at the protein level in participants with dandruff. These results strongly suggest that microbial dysbiosis contributes to dandruff pathogenesis through disruption of AhR signalling, a pathway crucial for maintaining epidermal barrier function and immune homeostasis. This *in vitro* model provides new insights into the role of microbial dysbiosis in skin disorders and could open new avenues for host-microbiome therapeutic interventions.

## Introduction

The skin is colonised by thousands of microbial species, including bacteria, fungi, viruses and mites, that make up the commensal microbiome^1^. The commensal microbiome has a vital role in skin barrier function, which is essential in maintaining skin health and homeostasis. In addition to the skin microbiome’s role in preventing colonisation by pathogenic microorganisms and priming the cutaneous immune system, the commensal microbiome has been shown to enhance epidermal barrier integrity through the control of keratinocyte proliferation, differentiation and cornification^1,2^.

Microbial dysbiosis has been associated with both acute and chronic skin conditions, contributing to pathogenesis by inducing skin barrier defects and proinflammatory responses^3^. An example of a skin condition commonly associated with microbial dysbiosis is dandruff, a chronic scalp condition that affects approximately 50% of the post-pubertal population and is characterised by visible flaking of the *stratum corneum* (SC), pruritus and mild inflammation^4–14^.

Several studies have shown that dysbiosis in dandruff is linked to five dominant skin commensals: *Staphylococcus epidermidis, Staphylococcus capitis, Cutibacterium acnes, Malassezia restricta* and *Malassezia globosa*. Specifically, in the mycobiome within dandruff, a significantly higher ratio of *M. restricta* to *M. globosa-* the two predominant fungal commensals are observed, correlating with increased symptom severity^6–9,15,16^. Similarly, in the core bacterial microbiome, a significant increase in the abundance of *Staphylococcus* species and a reduction in the relative abundance of *C. acnes* is observed^6–9,15,16^. Despite recognising the association between microbial dysbiosis and dandruff, the mechanism by which microbial dysbiosis develops and contributes to dandruff pathogenesis remains poorly understood. This is further highlighted by the limitations of traditional dandruff therapeutics, which primarily target the microbial population. Relapse of symptoms and further microbial dysbiotic changes are commonly observed following cessation of treatments^4,17–19^.

2D coculture models and mouse studies have advanced our understanding host-microbiome interactions in the skin, however several limitations of these systems impact the relevance of research outcomes^20–25^. 2D cell culture models do not possess the complex stratified structure of the epidermis that is an essential component of the skin as a protective layer. Mouse studies pose limitations due to differences in mouse and human skin physiology, biology and structure, as well as differences in the composition of the native microbiome population^26^.

3D skin organotypics (OTs) offer an easily adaptable, biologically representative *in vitro* tool for the investigation of the commensal microbiome. OTs form a stratified epidermis therefore, more effectively recapitulating host-microbiome interactions in the skin^27–29^. A small number of studies have documented the use of 3D skin OTs to investigate host-microbiome interactions in the skin; however, most of these models only involve single or dual species colonised OTs^30–34^, although a 3D skin OT with eight bacterial species of equal densities has been achieved. This model revealed that commensal microbes differentially influenced the epidermal barrier when in single species inoculations compared to a mixed community^33^. Despite these significant findings, improved models are required to better recapitulate the skin microbiome, specifically, to incorporate critical fungal species together with to bacterial species. Additionally, modelling the ratio of microbial species densities is essential to better model the observed ratios in the healthy and diseased human scalp skin. Such advances in 3D modelling will provide insight into interspecies interactions within the microbial population.

Here, microbially competent 3D skin OTs were developed using donor-matched normal human epidermal keratinocytes (NHEKs) and dermal fibroblasts (hDFs) isolated from healthy adult skin biopsies and colonised with five dominant members of the human skin microbiome: *S. epidermidis, S. capitis, C. acnes, M. restricta* and *M. globosa* (Fig. S1). A model healthy scalp microbiome inoculation, referred to as 5M, was developed and colonisation of OTs enabled investigation into the importance of a heterogenous microbiome population in skin health. A dandruff scalp microbiome, referred to as 5MP, was developed by altering the ratio of these microbes to model and investigate microbial dysbiosis in the pathogenesis of dandruff. The microbially competent 3D skin OT described here, not only enables detailed study of dandruff-associated dysbiosis, but also serves as a versatile tool for investigating the role of the microbiome in a range of other skin diseases.

## Results

### Modelling healthy scalp microbiome with 5M inoculation

To model the healthy scalp microbiome, an OT model was developed using NHEKs with a combined inoculation (5M) of *S. epidermidis, S. capitis, C. acnes, M. restricta* and *M. globosa* (Fig. S1). Single inoculations for each microbial species were based on an initial seeding density (CFU/ml) that caused the least cytotoxicity, as determined by LDH assay (Fig. S2A); whilst also enabling microbial colonisation to best mimic the host-microbiome relationship in healthy human skin (Fig. S2B). RT-qPCR was used to quantify the final abundance of each microbial species on OTs colonised with singles species inoculations and the 5M inoculation (Fig. 1A). This showed that the ratio of *S. epidermidis, S. capitis, C. acnes, M. restricta* and *M. globosa* was similar to ratios previously reported for healthy *in vivo* scalp skin, suggesting that 5M colonised OTs mimics healthy microbiome colonisation of human scalp skin (Fig. 1B)^6^.

**Figure 1.**
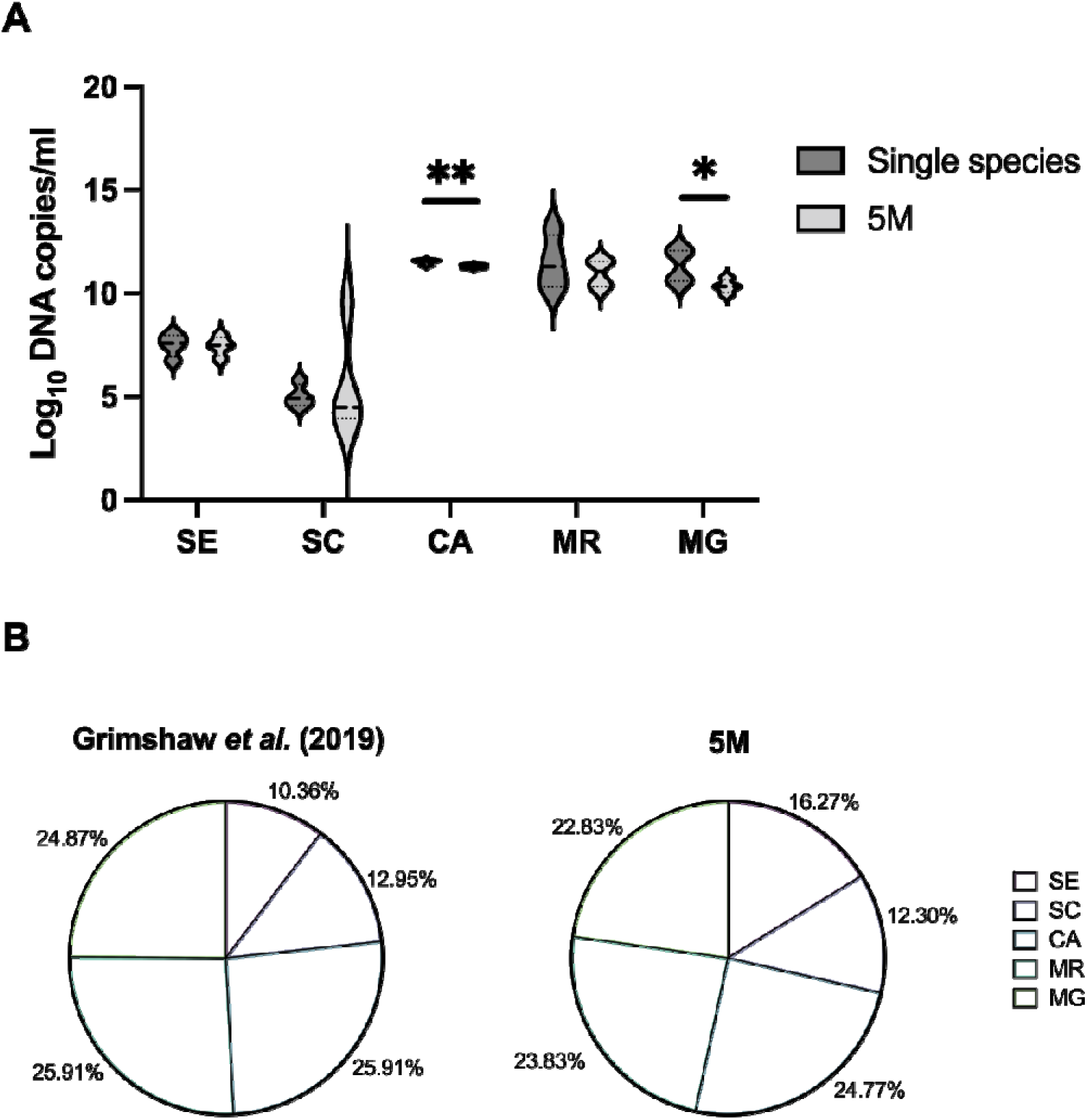
Quantification of the final abundance of microbes at harvest, following 3 days incubation on NHEK OTs, shows that 5M replicates the ratio of microbes observed on healthy scalp skin. (A) RT-qPCR analysis of each microbial species, represented as Log_10_ DNA copies/ml. Data represents n=4 biological replicates, carried out in triplicate, ± SD. Statistical analysis was carried out using separate paired t-tests for each microbe comparing quantity in single and 5M inoculations. *p≤0.05, **p≤0.01. (B) The ratio of *S. epidermidis*, *S. capitis, C. acnes, M. restricta* and *M. globosa* in healthy scalp microbiome sample analysis ^6^ compared to the 5M inoculation. *S. epidermidis* (SE), *S. capitis* (SC), *C. acnes* (CA), *M. restricta* (MR), *M. globosa* (MG), single species inoculations (singular) and combined microbe inoculation (5M).

Spongiosis was observed in OTs colonised by single species inoculations of *S. capitis*, *C. acnes, M. restricta* and *M. globosa* but not *S. epidermidis.* More disorganised epidermal stratification was observed in OTs colonised with single species inoculations of *C. acnes* and *M. globosa* (Fig. 2A). *S. epidermidis* only colonisation caused a significant decrease in epidermal thickness compared to sterile control OTs, from an average thickness of 50.6μm to 26.1μm (p=0.0032) (Fig. 2B). 5M colonisation showed a trend towards increased epidermal thickness, to an average of 66.7μm (p=0.2190) (Fig. 2B). All OTs exhibited a basal layer, indicated by K14 expression, and early epidermal differentiation, indicated by K10 in the suprabasal layers, similar to what is observed in healthy *ex vivo* skin (Fig. 2C, D and Fig. S3). Colonisation with these single inoculations of skin-resident microbes did not significantly influence K14 and K10 expression.

**Figure 2.**
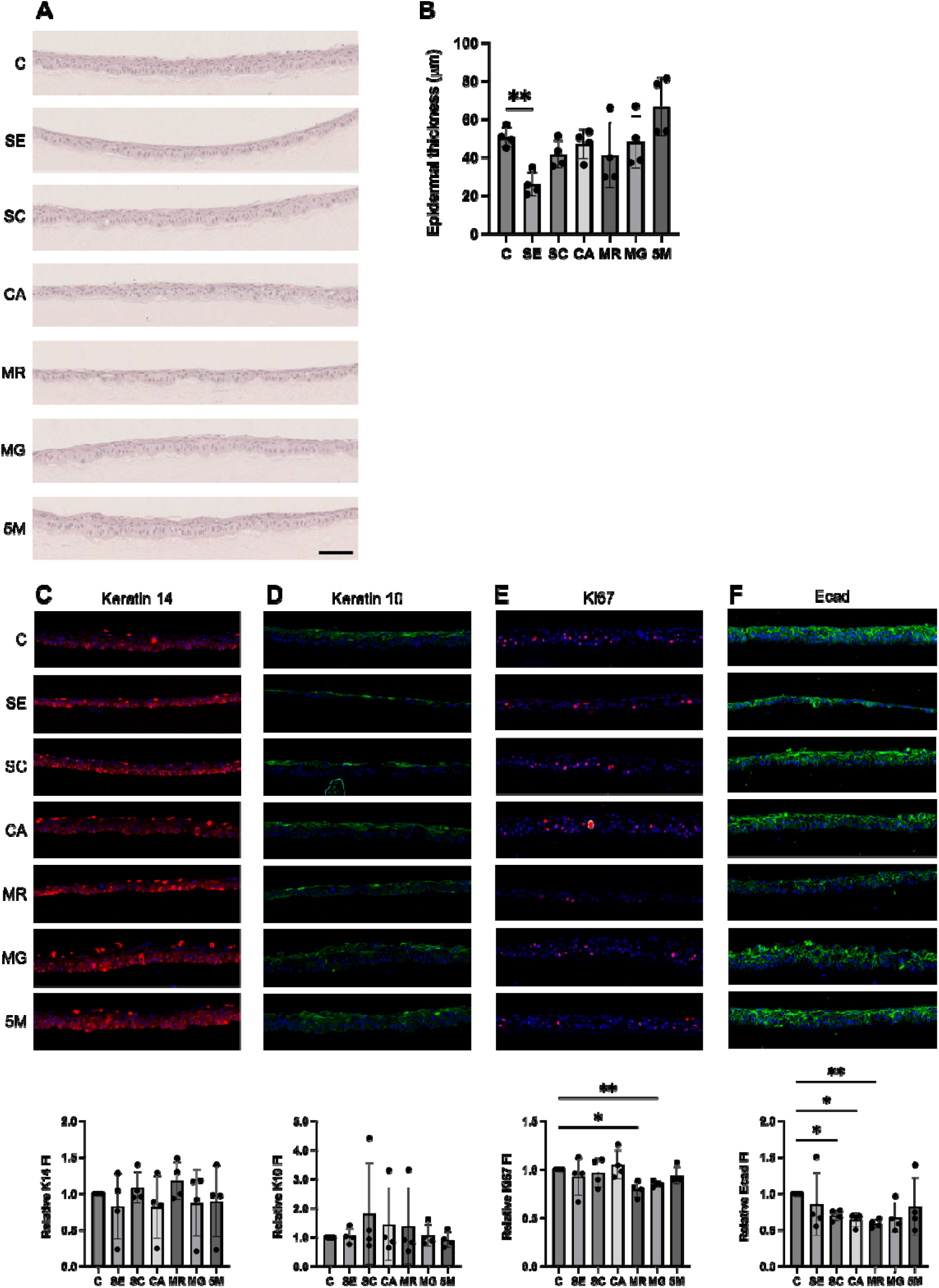
Morphological comparison of NHEK OTs colonised with *S. epidermidis, S. capitis*, *C. acnes*, *M. restricta*, *M. globosa*, or 5M compared to sterile control OTs. (A) Representative H&E images (Nanozoomer slide scanner, x20 magnification), scale bar = 250µm. (B) Epidermal thickness measurements taken from H&E images using NDP.view2 software (Hamamatsu Photonics). Representative IF images (INCA2200 widefield microscope, x20 magnification) and quantification (HALO software, Indica Labs) of fluorescent intensity (FI) of OTs immunolabelled for (C) Keratin 14 (K14, red channel), (D) Keratin 10 (K10, green channel), (E) Ki67 (red channel) and (F) E-cadherin (Ecad, green channel). Cell nuclei were visualised using DAPI (blue channel). Scale bar = 100µm. Sterile control (C), *S. epidermidis* (SE), *S. capitis* (SC), *C. acnes* (CA), *M. restricta* (MR), *M. globosa* (MG) and combined microbe inoculation (5M). Data represents n=4 biological replicates carried out in triplicate ± SD. Statistical analysis was carried out using one-way ANOVA with Dunnett’s multiple comparisons test. *p≤0.05, **p≤0.01.

A significant reduction in Ki67 (a marker of proliferating cells) expression was observed when OTs were colonised with *M. restricta* (p=0.0484) and *M. globosa* (p=0.0082), in comparison to sterile control OTs (Fig. 2E). E-cadherin (Ecad) is a cell-cell adhesion molecule expressed in the skin and was used as a marker of epidermal barrier function. A significant reduction in Ecad expression was observed in OTs colonised with *S. capitis* (p=0.0124), *C. acnes* (p=0.0146) and *M. restricta* (p=0.0028) (Fig. 2F).

In contrast to single species colonised OTs, 5M colonised OTs did not exhibit any significant changes in epidermal morphology and thickness (Fig. 2A-B), or changes in Ki67 and Ecad expression (Fig 2E-F) compared to sterile control OTs. Overall, 5M was more consistent with healthy *ex vivo* skin

### Developing a model of microbial dysbiosis

Several studies have demonstrated the association between dandruff and commensal microbiome dysbiosis^6,7,9,15,35,36^. To develop a model of microbial dysbiosis in dandruff the initial seeding density of *S. epidermidis, S. capitis, C. acnes, M. restricta* and *M. globosa* in the 5M inoculation was adjusted to model the average ratio observed in clinical studies of the dandruff scalp microbiome. RT-qPCR quantification of the final abundance of all five species on OTs colonised with 5MP compared to 5M confirmed a higher abundance of *S. epidermidis* and *M. restricta* and a lower abundance of *C. acnes* (p=0.0213) and *M. globosa* in 5MP, consistent with clinical dandruff microbiome analysis (Fig. 3A)^6,7,9,15^.

**Figure 3.**
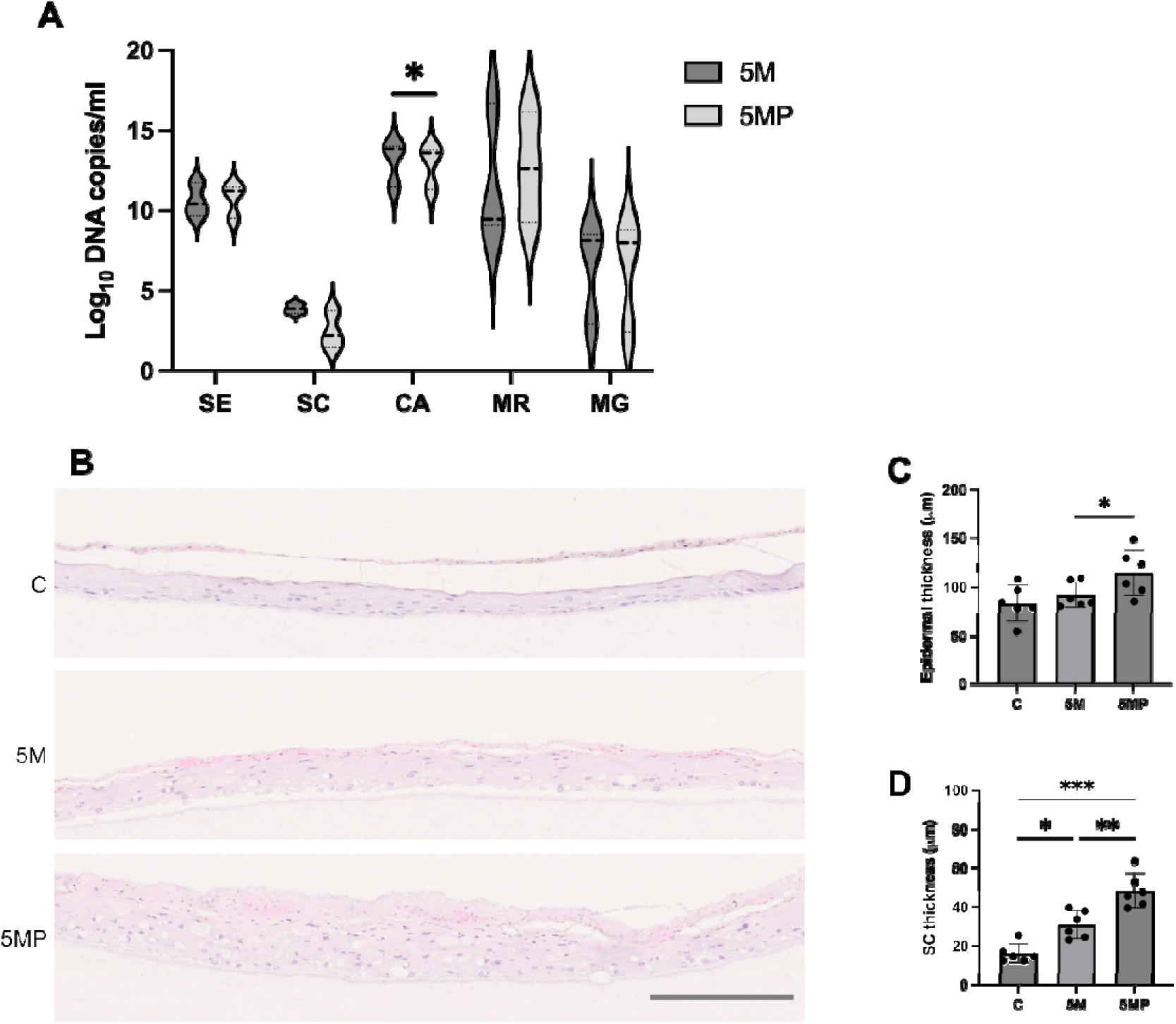
Colonisation with 5MP causes OTs to develop a dandruff-like phenotype, compared to 5M colonised OTs. (A) RT-qPCR analysis of each microbial species, represented as Log_10_ copies/ml, on OTs colonised with 5M or 5MP. Data represents n=3 biological replicates carried out in triplicate ± SD. Statistical analysis was carried out using separate paired t-tests. *p≤0.05. (B) Representative H&E images (Nanozoomer slide scanner, x20 magnification) of sterile control OTs, 5M colonised OTs colonised and 5MP colonised OTs, scale bar = 250μm. (C) Epidermal thickness measurements taken from H&E images using NDP.view2 software (Hamamatsu Photonics). (D) *Stratum corneum* thickness measurements taken from H&E images using NDP.view2 software (Hamamatsu Photonics). Data represents n=6 biological replicates carried out in duplicate ± SD. Sterile control (C), healthy microbiome inoculation (5M), dandruff microbiome inoculation (5MP). Statistical analysis was carried out using one-way ANOVA with Tukey’s multiple comparisons test. *p≤0.05, **p≤0.01, ***p≤0.001.

Colonisation of OTs with the altered 5MP microbiota resulted in the development of dandruff-like epidermal phenotypes akin to abnormal differentiation such as increased epidermal and SC thickness, spongiosis and increased parakeratosis in the *stratum corneum* (Fig. 3B). Furthermore, histological analysis revealed a significant increase in epidermal and *stratum corneum* thickness in OTs colonised with 5MP compared to 5M (Fig. 3B-D).

As with the previous OTs, all models exhibited early epidermal differentiations, indicated by the expression of K14 and K10 in the basal and suprabasal layers of the epidermis respectively (Fig. S4A, B). A significant reduction in the percentage of K10 positive cells was observed in 5MP colonised OTs compared to sterile control OTs (p=0.0357) (Fig. S4B). Although no significant differences in Ki67 expression was quantified, 5MP colonised OTs exhibited greater suprabasal expression of Ki67 (Fig. S4C).

### Microbial colonisation and dysbiosis influences gene expression in 3D skin OTs as revealed by bulk RNA-sequencing analysis

RNA-seq analysis was performed on sterile control OTs, 5M and 5MP colonised OTs to interrogate the effect of microbial dysbiosis at the transcriptional level and identify differentially expressed genes (DEGs). Principal component analysis revealed that OTs cluster based on treatment, with 5M and 5MP colonised OTs seperating from sterile control OTs, and distinct from each other (Fig. 4A). Differential expression analysis was carried out using Partek Flow using the DESeq2 statistical filter, where genes that had a significance value equivalent to or below 0.01 were considered differentially expressed. Of the 12,438 total mapped genes, 2,777 and 2,553 DEGs were identified when comparing 5M and 5MP colonised OTs respectively to sterile control OTs (Fig. 4B, C). When comparing 5MP colonised OTs to 5M colonised OTs, 160 DEGs were identified (Fig. 4B, C). The hierarchical clustering analysis of all DEGs identified between conditions shows that OTs cluster based on treatment, indicated as a heatmap plot (Fig. 4D).

**Figure 4.**
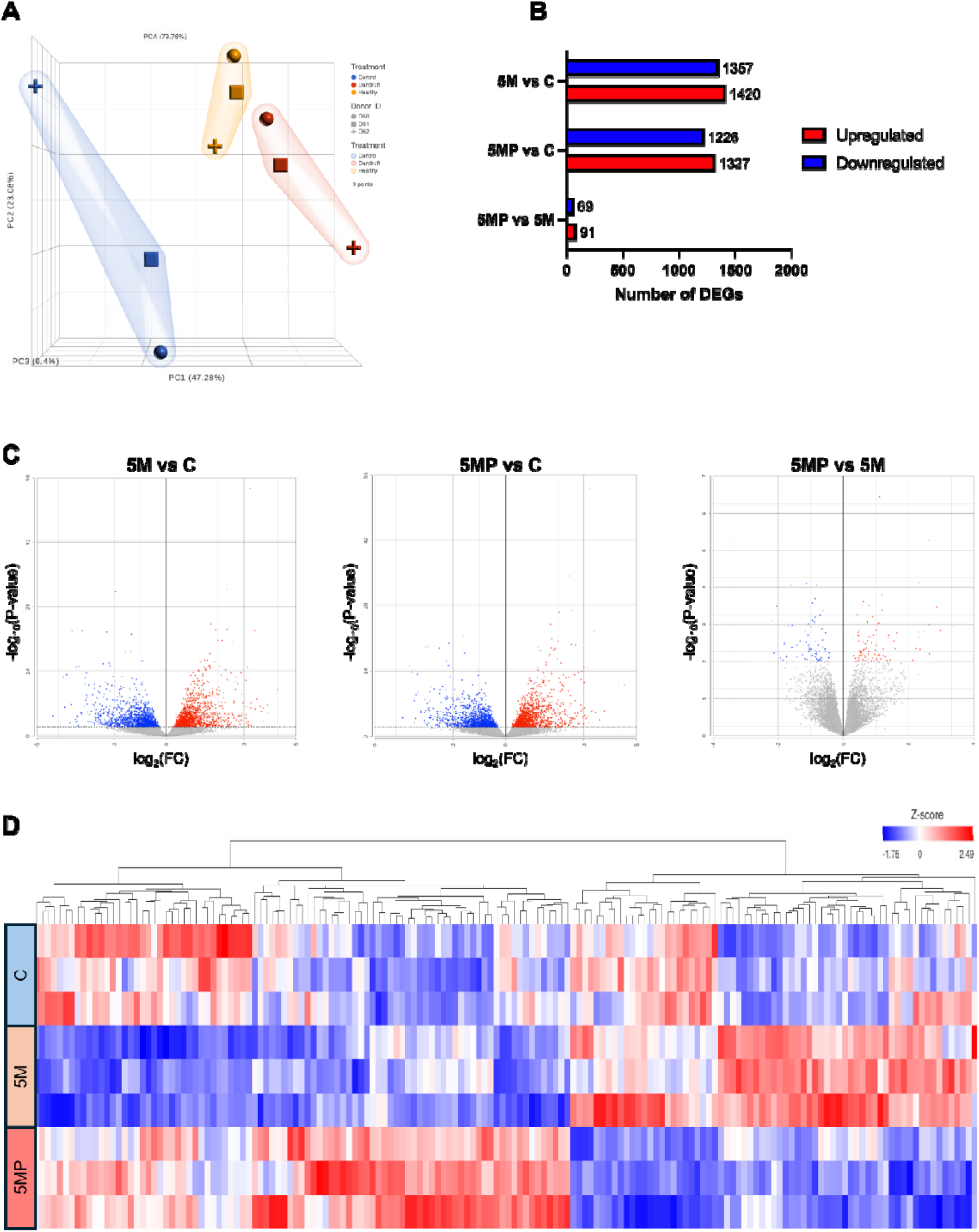
Transcriptomic analysis comparing sterile control OTs, 5M colonised OTs and 5MP colonised OTs. (A) PCA plot. (B) The number of differentially expressed genes (DEGs) detected. (C) Volcano plots showing log2FC and adjusted p values for genes. (D) Heatmap of significantly differentially expressed genes across all conditions, red indicates high expression values and blue indicates low expression values as z-scored normalised ratios. Significantly altered gene expression was based on an adjusted p value≤0.01. Sterile control (C), healthy microbiome inoculation (5M), dandruff microbiome inoculation (5MP). Data represents n=3 biological replicates.

### Dysregulated corneodesmosome hydrolysis in 5MP colonised OTs validates the development of dandruff-like phenotypes

Gene and protein expression analysis revealed changes in the expression of proteins involved in corneodesmosome hydrolysis, a process linked to increased SC thickening and flaking, in 5MP compared to 5M colonised OTs (Fig. 5). IPA was used to overlay gene expression data to model this prediction. Gene expression analysis revealed a 1.7-fold increase in *SPINK5* (encoding Lympho-epithelial Kazal-type-related inhibitor (LEKTI-1)) expression in 5MP colonised OTs, alongside a 1.1-fold reduction in *KLK5* expression and a predicted reduction in *KLK14* expression (Fig. 5A). Immunofluorescence was used to confirm these changes at the protein level. Increased levels of LEKTI-1 and SPINK6 and reduced levels of KLK5 were detected in 5MP colonised OTs, corroborating the gene expression analysis (Fig. 5B-D). A significant increase in LEKTI-1 and SPINK6 protein expression was observed in OTs colonised with 5MP compared to 5M, p=0.0372 and p=0.0400 respectively (Fig. 5B, C).

**Figure 5.**
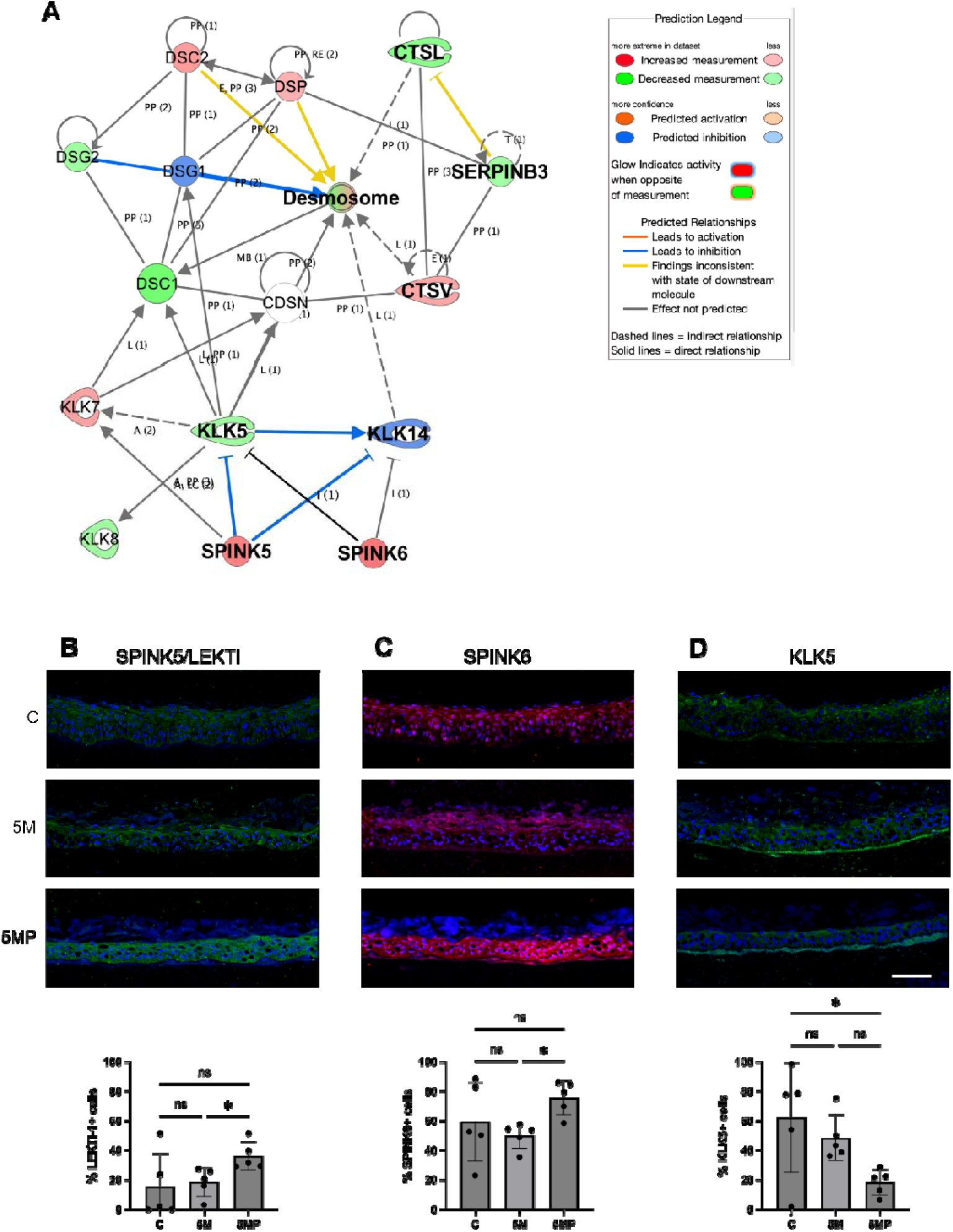
Transcriptomic analysis comparing 5M and 5MP colonised OTs suggests that microbial dysbiosis changes the expression of genes encoding proteins that regulate corneodesmosome hydrolysis. (A) Gene interaction network map overlayed with gene expression data using IPA (QIAGEN). Representative IF images (INCA2200 widefield microscope) of OTs immunolabelled for (B) LEKTI-1 (green channel), (C) SPINK6 (red channel) and (D) KLK5 (green channel) and quantification of percentage positive cells (HALO software, Indica Labs). Cell nuclei were visualised using DAPI (blue channel). Scale bar = 100µm. Sterile control (C), healthy microbiome inoculation (5M), dandruff microbiome inoculation (5MP). Data represents n=3 biological replicates carried out in duplicate ± SD. Statistical analysis was carried out using one-way ANOVA with Tukey’s multiple comparisons test. *p≤0.05, **p≤0.01.

### The AhR signalling pathway is attenuated in OTs colonised with 5MP compared to 5M

Gene expression analysis revealed enrichment of the AhR signalling pathway in response to microbial colonisation of 3D skin OTs. Comparative analysis revealed differential AhR signalling activity between 5M and 5MP colonised OTs, with a predicted inhibition of AhR signalling in 5MP compared to 5M (Fig. 6A). Microbial colonisation induced an increase in *AHR* expression in OTs, compared to sterile control OTs, with 5M and 5MP colonised OTs exhibiting a 1.9-fold and 1.7-fold increase in *AHR* expression respectively (Fig. 6B). However, when comparing 5MP to 5M colonised OTs, a 1.1-fold reduction in *AHR* expression was observed (Fig. 6B). Immunofluorescence showed that gene and protein expression of AhR corroborated. A significant increase in AhR expression was observed when comparing 5M colonised to sterile control OTs (p=0.0426) and a significant decrease in AhR expression was observed when comparing 5MP to 5M colonised OTs (p=0.0036) (Fig. 6C).

**Figure 6.**
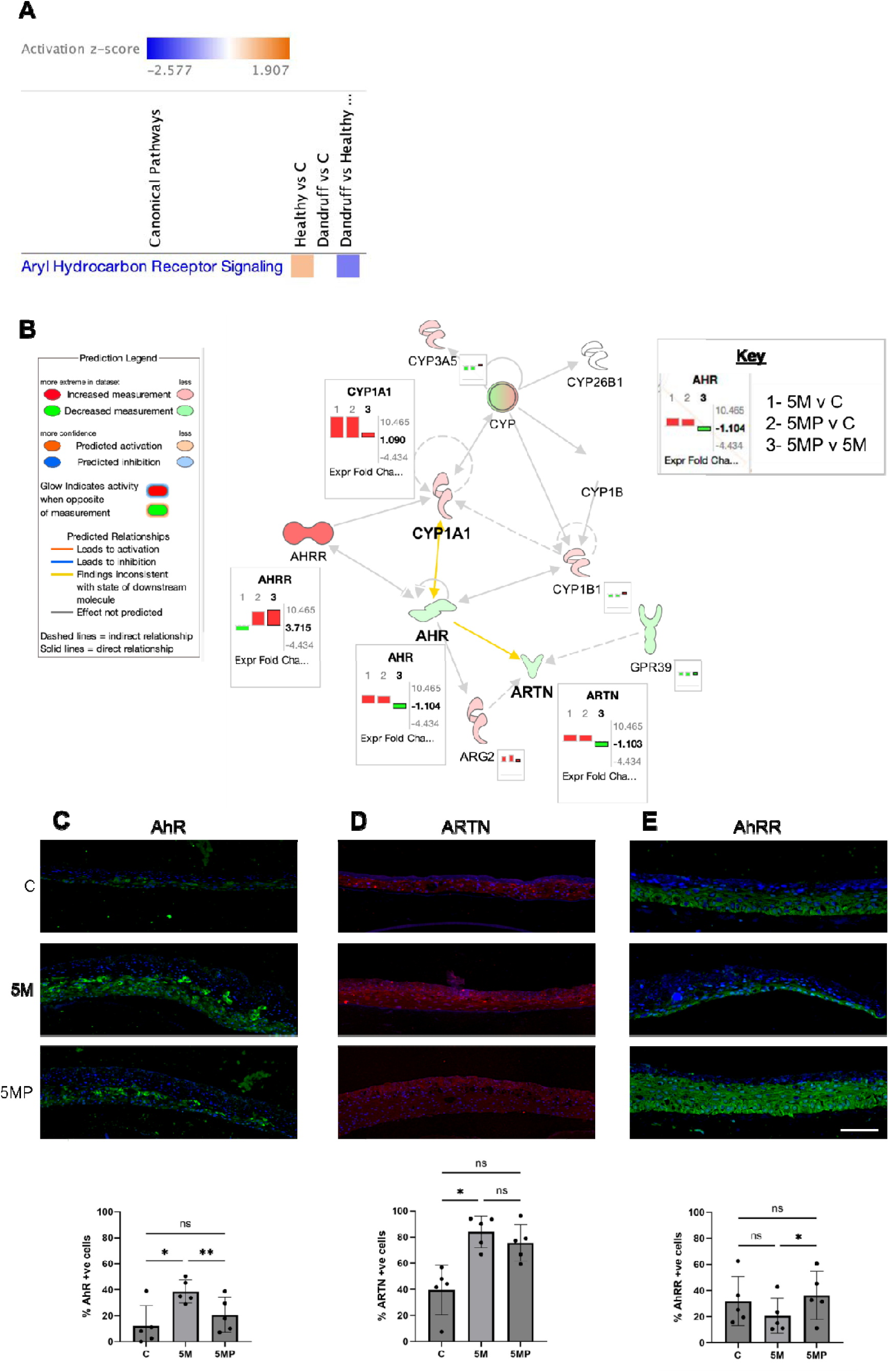
Transcriptomic analysis comparing 5M and 5MP colonised OTs suggests that microbial dysbiosis influences the AhR signalling pathway. (A) Comparison analysis predicted an inhibition of AhR signalling following 5MP colonisation compared to 5M colonisation. (B) Gene interaction network map of the AhR canonical signalling pathway overlayed with gene expression data using IPA (QIAGEN). Representative IF images (INCA2200 widefield microscope) of OTs immunolabelled for (C) AhR (green channel), (D) ARTN (red channel) and (E) AhRR (green channel) and quantification of percentage positive cells (HALO software, Indica Labs). Cell nuclei were visualised using DAPI (blue channel). Scale bar = 100µm. Sterile control (C), healthy microbiome inoculation (5M), dandruff microbiome inoculation (5MP). Data represents n=5 biological replicates carried out in duplicate ± SD. Statistical analysis was carried out using one-way ANOVA with Tukey’s multiple comparisons test. *p≤0.05, **p≤0.01.

Compared to sterile control OTs both 5M and 5MP colonised OTs exhibited an increase in ARTN gene and protein expression, corroborating with the AhR expression pattern observed (Fig. 6C, D). Interestingly, despite a reduction in AhR expression in 5MP colonised OTs, there was no significant change in ARTN expression, suggesting that microbial dysbiosis contributes to a dysregulated AhR-ARTN axis in 5MP colonised OTs.

The AhR signalling pathway is tightly regulated by several negative feedback loops, one of which is via the expression of the AhR repressor (AhRR)^37,38^. Gene expression analysis revealed that the different microbe inoculations distinctly influenced *AHRR* expression. A 1.1-fold reduction in *AHRR* gene expression was detected in 5M colonised OTs, whereas 5MP colonisation caused a 3.4-fold increase in *AHRR* gene expression, compared to sterile control OTs (Fig. 6B). When comparing 5MP to 5M, a 3.7-fold increase in *AHRR* gene expression was observed (Fig. 6B). This was validated at the protein level whereby AhRR expression trended towards a reduction when comparing 5M colonised to sterile control OTs and an increase in AhRR expression was observed when comparing 5MP colonised to 5M colonised OTs (p=0.0387) (Fig. 6E).

### Altered expression of AhR pathway proteins in lesional dandruff skin

AhR, ARTN and AhRR protein expression was assessed in healthy and lesional dandruff scalp biopsies to validate the observed dysregulation of the AhR pathway in 5M colonised OTs compared to 5MP colonised OTs (Fig. 7). Expression patterns of these proteins in adult healthy scalp skin compared to adult lesional dandruff skin corroborated with the gene expression and protein expression data generated from the colonised 3D skin OTs (Fig. 6). Lesional dandruff skin exhibited a reduction in AhR protein expression when compared to healthy control skin, akin to what was observed in the colonised OTs (Fig. 6C and Fig. 7A, B). Similarly, the increased expression of AhRR in lesional dandruff compared to healthy control skin, corroborated the 3D skin OT analysis (Fig. 6E and Fig. 7E, F).

**Figure 7.**
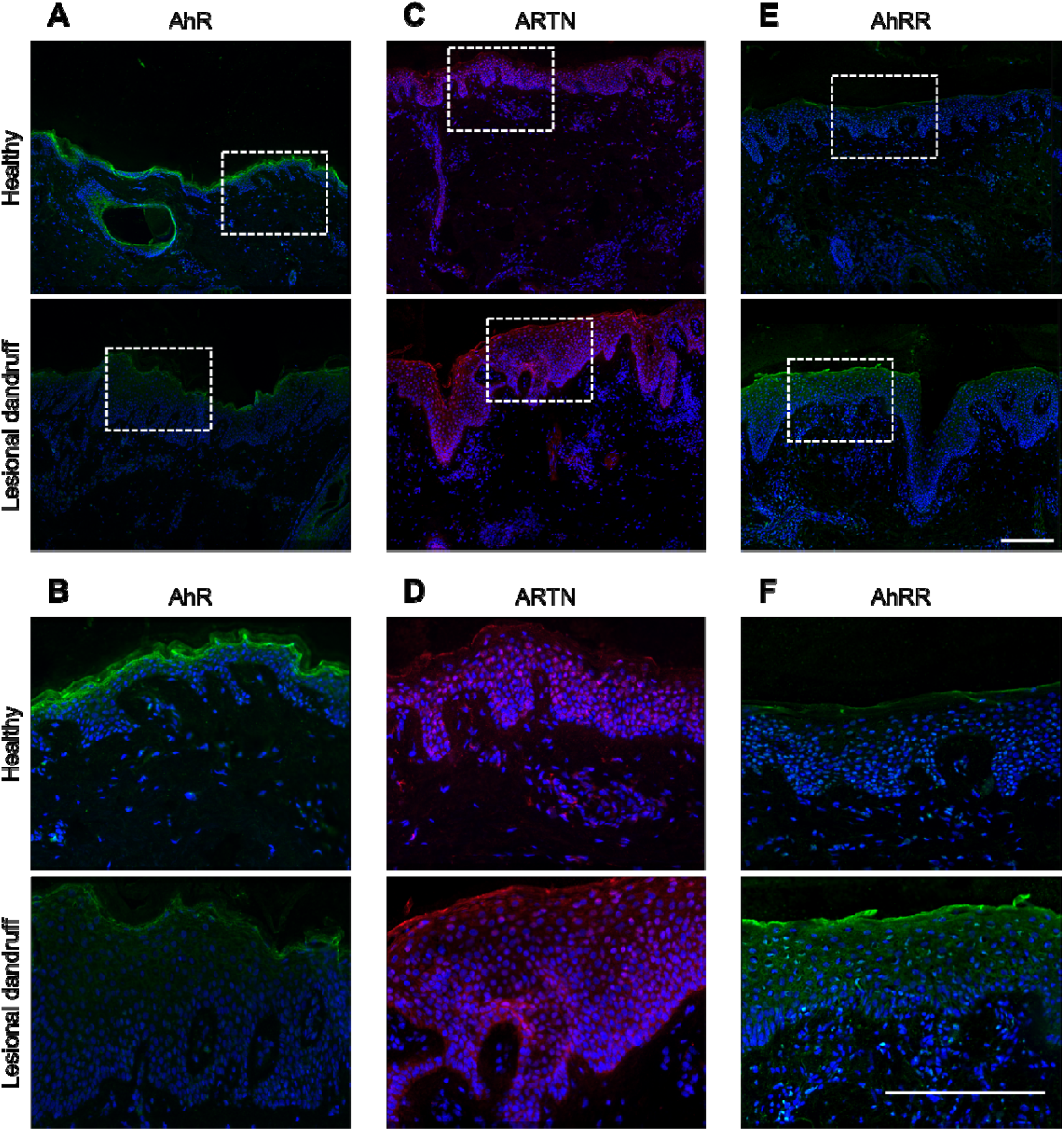
Expression of AhR, ARTN and AhRR in human adult lesional dandruff skin compared to human adult healthy skin. Representative IF images (INCA2200 widefield microscope) of human adult lesional dandruff and healthy control skin immunolabelled for (A) AhR (green channel), (B) magnified area of (A) indicated by white box, (C) ARTN (red channel), (D) magnified area of (C) indicated by white box, (E) AhRR (green channel) and (F) magnified area of (E) indicated by white box. Cell nuclei were visualised using DAPI (blue channel). Data represents n=5 biological replicates. Scale bars = 200µm.

Finally, despite a clear difference in AhR and AhRR protein expression when comparing lesional dandruff and healthy control skin, overall, no change in ARTN expression was observed (Fig. 7C, D). This was also observed when comparing 5MP and 5M colonised OTs (Fig. 6D). This could further suggest that the AhR-ARTN axis is dysregulated in response to a dysbiotic skin microbiome.

## Discussion

Here, microbially competent 3D skin OTs were developed to investigate how the scalp microbiome influence epidermal homeostasis and to examine the impact of microbial dysbiosis on host responses. Initially, microbial colonisation of OTs using single species inoculations was employed to investigate how individual members of the microbiome may influence the skin. The 5M inoculation was developed to model the healthy skin microbiome, enabling investigation into the importance of a heterogenous microbiome population in skin health. Finally, the 5MP inoculation was developed to model microbial dysbiosis in dandruff, and its colonisation of OTs was used to investigate how microbial dysbiosis influences skin health. Using these OTs, attenuation of the AhR pathway was identified as a mechanism by which microbial dysbiosis may contribute to the pathogenesis and persistence of dandruff.

OTs colonised with 5M exhibited an epidermal morphology most similar to *in vivo* skin. (53)(32). By quantifying the final abundance of each microbe on 5M colonised OTs and comparing to published data on the healthy scalp microbiome, similar ratios of each microbe in the 5M inoculation were observed^6^. A conversion factor for each species was calculated, enabling the adaptation of initial seeding densities of 5M to generate 5MP which represented the dandruff microbiome^6^. RT-qPCR confirmed the 5MP as a suitable *in vitro* model of the dandruff microbiome when the final abundance of microbes on OTs colonised with 5M and 5MP were compared.

5MP colonisation led to the development of dandruff-like phenotypes in OTs, compared to 5M. Histological analysis revealed significant changes in epidermal morphology with OTs exhibiting increased epidermal and SC thickness, parakeratosis and spongiosis, recapitulating what is observed in dandruff skin^5,11,39^. 5MP colonised OTs showed a significant reduction in the expression of the early epidermal differentiation protein K10, which has been observed in dandruff and other skin conditions associated with epidermal barrier dysfunction, including psoriasis and AD^5,40,41^.

In dandruff skin, the presence of flakes and increased SC thickness has been attributed to abnormal corneodesmosome retention and desquamation^10,11^. For desquamation to occur, corneodesmosomes are hydrolysed by serine proteases, such as kallikreins and cathepsins, in a process that is regulated by serine protease inhibitors such as LEKTI-1 and SPINK6. Despite the inability to model desquamation *in vitro*, visual assessment showed morphological changes in the SC of 5MP colonised OTs indicated abnormal corneodesmosome hydrolysis, as observed in dandruff skin^10–13^.

Pathway analysis predicted a reduction in the activity of the corneodesmosome proteases, KLK5 and KLK14, likely due to increased expression of genes encoding the protease inhibitors *SPINK5* and *SPINK6*, an observation consistent with what is observed in dandruff skin^10,42^. Immunofluorescence analysis of the expression of LEKTI-1 (encoded by *SPINK5)*, SPINK6 and KLK5 validated the gene expression changes at the protein level. Thus, suggesting that microbial dysbiosis is influencing corneodesmosome hydrolysis in dandruff. It is important to note that commensal microbes secrete proteases that contribute to the process of corneodesmosome hydrolysis in the skin, therefore differences in 5MP and 5M responses may be due to their protease secretome^43^.

The aryl hydrocarbon receptor (AhR) is a ligand-activated transcription factor that has a significant role in regulating epidermal barrier function and repair through the expression of xenobiotic metabolising enzymes (predominantly CYP1 enzymes) and proteins of the epidermal differentiation complex (EDC) (Fig. S5). Recently, the role of the commensal microbiome in mediating AhR signalling in the skin has been elucidated^44–47^. However, despite its beneficial functions, dysregulated AhR signalling is implicated in several chronic inflammatory skin conditions such as atopic dermatitis and psoriasis. Studies show that mice deficient in epithelial AhR have an impaired epidermal barrier, indicated by increased TEWL and reduced expression of epidermal differentiation genes such as *KRT10* and *IVL*^46,48^. Additionally, it has been demonstrated that AhR deficiency in imiquimod-induced psoriasiform mouse models exacerbates IL-17 and IL-22 mediated skin inflammation, highlighting the immunosuppressive role of AhR signalling in mediating immune responses^45^. The beneficial effects of AhR activation in the skin are also supported by the clinical use of AhR agonists, such as coal tar and Tapinarof, in the treatment of inflammatory skin conditions which have been shown to ameliorate disease symptoms by inducing AhR-dependent epidermal repair^49–52^.

Consistent with published reports of microbial colonisation inducing *AHR* expression in germ-free (GF) mice, an increase in AhR expression and signalling was observed in microbially colonised OTs compared to sterile control OTs (Fig. 6)^46^. Uberoi *et al.* show that colonisation of GF mice with ‘Flower’s Flora’ restores epidermal barrier function and the expression of EDC proteins via the AhR signalling pathway^46^. These findings collectively highlight the intricate relationship between the commensal microbiome, AhR, and skin homoeostasis.

The AhR pathway is controlled by self-regulating negative feedback loops is the AhR/ARNT-dependent transcriptional induction of *AHRR* expression^37,53^. While the exact mechanisms of AhRR-mediated AhR repression remain to be elucidated, studies suggest that AhRR competes with AhR for ARNT dimerization and XRE-containing gene promoter binding, thus inhibiting nuclear translocation and transcriptional activity^37,54,55^. A significant increase in AhRR expression, at the mRNA and protein level, has been observed in psoriasis skin compared to healthy control skin due to DNA hypomethylation of the *AHRR* gene^56^.

Strikingly, a 3.7-fold increase in *AHRR* expression, and a significant increase in AhRR protein expression was observed when comparing 5MP colonised OTs to 5M colonised OTs. Thus, microbial dysbiosis in dandruff may attenuate AhR signalling through the regulation of AhRR expression.

Dysregulated or attenuated AhR pathway signalling has been implicated in several inflammatory skin conditions, including psoriasis and atopic dermatitis, but not previously reported in dandruff ^38,45,46,49–52,57^. Human skin analysis comparing healthy scalp skin, and lesional dandruff scalp skin showed lower levels of AhR and higher levels of AhRR were expressed in lesional dandruff skin. These findings corroborate and validate the results of the colonised OTs.

The expression of the neurotrophic factor artemin (ARTN) is mediated through the activation of the AhR signalling pathway^58–60^. ARTN has been shown to cause alloknesis (hypersensitivity to itch) by stimulating the elongation of peripheral nerve fibres into the epidermis^59–61^. Additionally, a dysregulated AhR-ARTN axis has been reported in contact dermatitis and AD patients^60^. The observed conflicting ARTN expression levels in lesional dandruff skin suggest a potentially dysregulated AhR-ARTN axis, as hypothesised in the OTs. Notably, this analysis reveals that lesional dandruff skin exhibits altered levels of proteins involved in the AhR signalling pathway, which may be linked to the characteristic microbiome dysbiosis of dandruff. Combining these findings with previously published data it is plausible to hypothesise that the dysbiotic microbiome in dandruff contributes to altered AhR expression and signalling.

This study has provided valuable insights into the relationship between the skin microbiome and epidermal homeostasis in the context of dandruff. By developing microbially competent 3D skin OTs, we have demonstrated how dandruff-associated microbial dysbiosis leads to alterations in epidermal morphology, barrier function, and molecular processes. The attenuation of the AhR pathway in the context of microbial dysbiosis represents an understudied mechanism that may contribute to the development and persistence of dandruff. Continuing to unravel the complex relationship between the commensal microbiome, epidermal homeostasis, and key signalling pathways, such as the AhR, will enable the development of more targeted and effective treatments for dandruff and other skin conditions associated with microbial dysbiosis.

## Materials and Methods

### Human adult healthy scalp skin from elective rhytidectomy surgery

Healthy adult human skin was obtained from redundant skin tissue with written and informed consent from participants undergoing elective rhytidectomy surgery (LREC No. 09/HO704/69).

### Human adult lesional dandruff and healthy control scalp skin samples

Human scalp specimens were obtained from healthy (eight subjects) and dandruff subjects (eight subjects) by Alba Science Limited (Edinburgh) on behalf of Unilever PLC under ethical approval and written informed subject consent. Ethical approval was granted by the Reading Independent Ethics Committee (Ethics Approval HAI-HUB-3577).

### Human primary keratinocyte and fibroblast isolation from normal human skin samples

Normal human skin samples were incubated in dispase II (2.5 mg/ml) overnight at 4°C to separate the epidermis and dermis. The epidermis was incubated in trypsin to isolate NHEKs and the dermis was incubated in trypsin for 2hrs, followed by collagenase D (0.5 mg/ml) overnight at 37°C to isolate hDFs.

### Mammalian cell culture conditions

Cell culture was performed in a laminar flow hood under aseptic conditions. All sterile disposable tissue culture flasks, plates and dishes were purchased from Nunc (Roskilde, Denmark) and all centrifuge steps were performed using an IEC Centra-3C Centrifuge (International Equipment company, Dunstable, UK). Cells and 3D skin organotypics were incubated at 37°C in a cell culture incubator at 5% CO_2_/ 95% atmospheric air.

Normal human epidermal keratinocytes (NHEKs) and the immortalised N/TERT keratinocyte cell line (NTERTs) were cultured in RM+ media (Dulbecco’s Modified Eagle’s Medium (DMEM)/F12 + GlutaMAX™ supplement, Gibco, Paisley, U.K.) with 10% (v/v) foetal bovine serum (FBS, Biowest, East Sussex, UK), 100 U/ml penicillin, 100 mg/ml streptomycin (Sigma-Aldrich, Poole, U.K.) supplemented with 5μg/ml apo-transferrin (Sigma-Aldrich, Poole, U.K.), 1×10^−8^M cholera toxin (Sigma-Aldrich, Poole, U.K.), 10ng/ml epidermal growth factor (Sigma-Aldrich, Poole, U.K.), 0.4μg/ml hydrocortisone (Sigma-Aldrich, Poole, U.K.), 5μg/ml insulin (Sigma-Aldrich, Poole, U.K.) and 2×10^−11^M 3,3’-5’-triiodo-L-thyronine (Sigma-Aldrich, Poole, U.K.). NHEK stocks were expanded with 10μM ROCKi (Y-27632, Bio-Techne Ltd, Abingdon, U.K.) and at least 12-16h prior to passaging for experiments, complete RM+ containing no ROCKi was used to ensure a sufficient wash-out period.

Normal human dermal fibroblasts (hDFs) were cultured in High-Glucose DMEM + GlutaMAX™ supplement (Gibco, Paisley, U.K.) supplemented with 10% (v/v) FBS, 100 U/ml penicillin and 100 mg/ml streptomycin.

### Microbial cell culture conditions

Frozen stocks of microorganisms were stored long term at -80°C and defrosted for culture when required. All microbial strains used in this study were kindly provided by Sally Grimshaw (Unilever R&D, Port Sunlight, UK). The strains were: *Staphylococcus epidermidis* NCTC 12228*, Staphylococcus capitis* from healthy skin isolate*, Cutibacterium acnes* NCTC 737*, Malassezia restricta* from healthy skin isolate and *Malassezia globosa* from healthy skin isolate. Culture conditions used for each microbial species are detailed in Table S1.

All aerobes were cultured in an aerobic atmosphere containing 5% CO_2_/95% atmospheric air. *Staphylococcus* species were cultured at 37°C and *Malassezia* species were cultured at 32°C. *C. acnes,* a facultative anaerobe, was cultured in a Whitely A35 Anaerobic Workstation (Don Whitely Scientific Ltd, Shipley, UK) with an atmosphere containing 80% N_2_, 10% H_2_ and 10% CO_2_ and maintained at 37°C. 24 hours prior to experiments, a single colony from each maintained culture was selected and resuspended in the appropriate broth and cultured overnight. In coculture with human cells, all cultures were incubated at 37°C in an incubator at 5% CO_2_/95% atmospheric air.

### OD_600_ and CFU/ml standard calibration curves

**Log-phase** batch cultures were adjusted to an OD_600_ of 0.1 for *S. epidermidis, S. capitis* and *C. acnes* and an OD_600_ of 1.0 for *M. restricta* and *M. globosa*, using the Eppendorf BioPhotometer Spectrophotometer Model #6131 (Eppendorf, Hamburg, Germany). Serial decimal dilutions of the microbial suspensions were spread on the appropriate agar, and colonies counted after an appropriate period of incubation to generate the standard calibration curves. GraphPad Prism (Version 10.1.1, San Diego, California, U.S.A.) was used to calculate linear regression to relate CFU/ml to the culture OD (Fig. S6).

### 2D NTERT coculture with serially diluted microbes

All coculture experiments were performed in RM+ cell culture media without antibiotics. NTERTs were seeded at a density of 1×10^5^ cells per well of a 12-well plate and cultured for 24hrs prior to microbial exposure.

Microbial suspensions were prepared from overnight broth cultures by pelleting at 2400 x g for 10 mins at 10°C and washing twice with 1XPBS before resuspending in RM+ culture media. A batch culture of each microbe was adjusted to an OD_600_ of 0.1, and serially diluted (1:10) to generate dose response curves. NTERTs were cocultured with 1ml from each microbe dilution for 3 days in an aerobic atmosphere containing 5% CO_2_ at 37°C.

Daily media changes were performed to remove planktonic microbes and encourage biofilm formation. On the final media change, RM+ media without FBS was used (as FBS is known to interfere with the LDH assay). After 36 hrs of coculture, an LDH colormetric assay (Roche) was performed on cell culture supernatants to assess cytotoxicity. NTERTs used for positive controls were incubated in a Triton X-100 solution (2% (v/v) in RM+) for 2 hrs, prior to supernatant harvesting.

Seeding densities were converted from OD_600_ to CFU/ml using standard calibration curves (Fig. S6).

### Lactate Dehydrogenase (LDH) assay

The LDH assay was performed using the Cytotoxicity Detection Kit (Roche, Welwyn Garden City, U.K.) according to manufacturer’s instructions. Cell culture supernatants were harvested and centrifuged at 250 *g* for 10 mins to pellet any cellular debris. 100ml of supernatant (avoiding the pellet) was transferred to a 96-well plate (VWR, Poole, U.K.) and 100ml of LDH reaction mix (250ml of catalyst per 11.25ml of dye) was added to the supernatant. The plate was then incubated for 30 mins protected from light at RT. Following incubation, the plate was read at 492nm on a CLARIOstar plate reader (BMG Labtech, Ortenberg, Germany). Percentage cytotoxicity was calculated using the following equation:

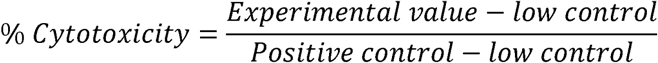

Positive control samples were supernatants collected from NTERTs that had been exposed to 2% (v/v) Triton X-100 for 2 hrs and low controls were sterile control NTERTs cultured in RM+.

### 3D skin organotypic generation

3D skin organotypics (OTs) were cultured in 12 well inserts with a PET membrane, pore size: 0.4μm (Sarstedt Ltd, Nümbrecht, Germany) in 12 well plates (VWR, Poole, U.K.). OTs were cultured in RM+ media and media was changed daily for the duration of culture. OT gels were prepared by combining 177.40 µl of collagen I (4 mg/ml), 66.60 µl of matrigel, 34 µl 10X MEM, 34 µl FBS and 34 µl of hDFs resuspended at 5×10^4^ in DMEM and seeded in a 12-well plate transwell insert. Following even resuspension of hDFs in the collagen-matrigel solution, 350μl of the gel matrix was transferred into each insert. Polymerisation gel matrix was promoted by incubating at 37°C for one hour. After the collagen gel was set, 2ml of DMEM was added into the well and 0.5ml was added on top of the transwell to ensure the gel was fully submerged for overnight incubation. The following day all media above and below the dermal matrix was removed. 1×10^6^ NHEKs in 500μl of RM+ culture media was seeded on top of the dermal matrix and 1ml of RM+ culture media was added underneath the insert. OTs were cultured submerged for 3 days, with daily RM+ media changes above and below the insert. On day 4 of culture, OTs were raised to the air-liquid interface on day 4 and grown for a further 10 days with daily RM+ media changes.

### Microbial colonisation of 3D skin organotypics

On day ten of culture, media was replaced with antibiotic-free RM+ media in preparation for microbe inoculation and used for the remainder of the experiment. The following day, broth cultures of each microbial species were centrifuged for 10 mins at 3500rpm and resuspended in 2ml of 1X PBS. The concentration of each microbial species was adjusted to the optimised OD_600_ measurements using the Eppendorf BioPhotometer Spectrophotometer Model #6131 (Eppendorf, Hamburg, Germany). Inoculated OTs were incubated at 37°C for 3 hours to encourage microbes to adhere to the surface of OTs. After 3 hours, any remaining PBS on the surface of the OT was removed to maintain the air-liquid interface. 250μl of sterile 1X PBS was added to the surface of sterile control OTs. OTs were cultured for a further 3 days until day 14 of culture, with daily antibiotic-free RM+ media changes. At the end of the experiment, OTs were removed from cell culture inserts and cut in half using a sterile scalpel. One half of the OT was fixed in 4% paraformaldehyde in PBS (4% PFA) for 1 hour and then stored in 70% ethanol for tissue processing. The other half of the OT was either snap frozen and stored at - 80°C for DNA extraction or submerged in RNA*later*™ and stored at -80°C for RNA extraction.

### Immunofluorescence and histological analysis of Ots

Formalin-fixed paraffin-embedded OTs were sectioned at 5µm on a microtome, transferred to SuperFrost Plus Adhesion slides and deparaffinised prior to staining. For histology, sections were stained using haematoxylin and eosin (H&E) and imaged using a Nanozoomer Slide Scanner. Deparaffinised sections underwent permeabilization with 0.1% Triton X-100 in 1X PBS and antigen retrieval using 1X citrate buffer (pH 6.0) or 1X Tris-EDTA (pH 9.0) prior to immunolabelling with antibodies (Table S2). Immunolabelled sections were imaged using an INCA 2200 widefield microscope (GE Healthcare) at 40x magnification. Analysis and IF quantification were performed using HALO software (Indica Labs).

### Immunofluorescence and histological analysis of human adult skin samples

Frozen OCT-embedded human adult skin was sectioned at 5µm on a cryostat, transferred to SuperFrost Plus Adhesion slides and fixed in 4% PFA for 10 mins prior to staining. For histology, sections were stained using haematoxylin and eosin (H&E) and imaged using a Nanozoomer Slide Scanner. For immunofluorescence analysis, sections were permeabilised with 0.1% Triton X-100 in 1X PBS prior to immunolabelling with antibodies (Table S2). Immunolabelled sections were imaged using an INCA 2200 widefield microscope (GE Healthcare) at 20x magnification.

### DNA extraction from 3D skin Ots

Snap frozen halves of models were suspended in 200µl of cetyltrimethylammonium bromide (CTAB) lysis buffer (AppliChem), 1µl of proteinase K (20mg/ml, Qiagen) and 1µl of Lysozyme (50mg/ml, Thermofisher) and incubated at 65°C for 3 hours to promote cell lysis. The lysis solution was transferred to a 2ml Lysing Matrix E (MP Biomedicals) tube with an additional 800µl of CTAB lysis buffer and homogenized using a TissueLyser LT (Qiagen) at 50Hz for 10 mins. Cell lysates were transferred to 1.5ml centrifuge tubes, incubated at 65°C for 1 hr and centrifuged at 12,000 g for 10 mins at 4°C and supernatants were transferred to fresh 1.5ml centrifuge tubes. 700µl of UltraPure™ Phenol: Chloroform: Isoamyl alcohol (P: C: I, 25:24:1, v/v, Invitrogen) was added to each sample, vortexed and incubated at RT for 10 mins. Centrifugation at 12,000 g for 10 mins at 4°C caused phase separation and the upper aqueous layer was transferred to a fresh 1.5ml centrifuge tube, this was repeated twice more with the final extraction step using 700µl of chloroform (Sigma): isoamyl alcohol (Sigma) (24:1, v/v) only to reduce phenol contamination. To precipitate DNA, 450µl of isopropanol (Sigma) and 75µl of 5M ammonium acetate (Invitrogen) was added to the samples overnight at -20°C. The following day samples were centrifuged at 12,000 g for 10 mins at 4°C and the supernatants were discarded. The DNA pellets were washed twice in 100µl of 80% (v/v) ethanol (Thermofisher) and the final pellet was air-dried for 5 mins and resuspended in 20µl of 1X Tris-EDTA (TE) buffer (pH 8.0, Thermofisher). DNA quality and concentration was analysed using the NanoDrop 8000 spectrophotometer (ThermoScientific) and DNA samples were stored at - 20°C until required.

### Real time-quantitative polymerase chain reaction (RT-qPCR)

qPCR amplification primers (Table S3) and probes (Table S4) (TaqMan) were designed based on previously published sequences. qPCR reactions were performed in MicroAmp™ Optical 96-well reaction plates (Applied Biosystems™) with a total reaction volume of 20µl. The qPCR reactions contained 10µl of Premix Ex Taq™ (Takara), 2µl of primer mix (containing 5µM of forward primer and 5µM of reverse primer), 2µl of 10µM probe, 2µl of template DNA and 4µl of nuclease-free water. qPCR plates were sealed with MicroAmp™ Optical Adhesive Film (Applied Biosystems™) and run on a StepOnePlus™ Real-Time PCR System (Applied Biosystems™). The amplification conditions were as follows: initial denaturation at 94°C for 2 mins, followed by 50 cycles of incubation at 94°C for 15s and annealing at 60°C for 1 min. Data was collected and Ct values calculated using the StepOnePlus™ software.

### RT-qPCR standard curves

Standard curves were generated to determine the quantity of genomic DNA of bacteria and fungi colonising tissue and OTs using Gene Fragments by GENEWIZ (Azenta, Germany). Gene Fragments are double-stranded DNA fragments of 100-3,000bp of known stock concentration (confirmed by Nanodrop) that can be serial diluted to amplify different amounts of the same sample to generate standard curves of Log_10_ amplicon copy number vs Ct values. Three Gene Fragments were generated: one for each genus of microbe. Each Gene Fragment contains forward and reverse primer sequences for each primer pair of the appropriate microbe. Table S5 details the gene fragment sequences used.

Seven serial dilutions were made from each sample of stock DNA (of known concentration), and a qPCR reaction was performed to generate the corresponding Ct values. qPCR reactions were performed in triplicate, and the mean Ct value was used to generate standard curves against Log_10_ DNA copy number (Fig. S7).

To calculate DNA copy numbers, the following equation was used:

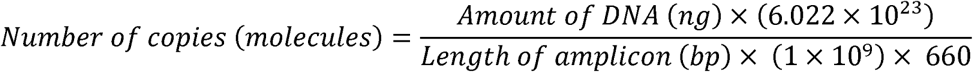

Whereby; 6.022×10^23^ is Avogadro’s constant (mol^−1^) which is used as a normalisation factor and represents the number of molecules in one mole of any substance, 660 (g/mol) represents the average mass of 1bp of dsDNA and 1×10^9^ is a conversion factor.

### RNA extraction

Total RNA was extracted from OTs using the RNeasy Lipid Tissue Mini Kit (QIAGEN, Hilden, Germany) following manufacturer’s instructions. RNA samples were assessed for quantity and integrity using an Agilent 2100 Bioanalyzer (Agilent Technologies).

### RNA-seq analysis

RNA samples were sent to Lexogen (Vienna, Austria) at a concentration of 50ng/µl in 20µl of nuclease free water for quality control, library preparation and sequencing. The QuantSeq 3’ mRNA Library Prep Kit FWD (Lexogen, Vienna, Austria) was used to generate next generation sequencing library sequences close to the 3’ end of polyadenylated RNAs. cDNA libraries were sequenced with an Illumina NextSeq2000 platform. The QuantSeq Data Analysis Pipeline was performed on FASTQ files by Lexogen prior to data delivery: Sequencing quality control of the raw reads was assessed using FastqQC software and adapter sequences were removed with cutadapt^62^. Alignment to the human reference genome (hsa_GRCh38) and read counting were performed using STAR and featureCounts ^63,64^. Batch effect removal and gene specific analysis was performed using *Partek™ Flow™* software, v11.0. (Partek Incorporated, Missouri, U.S.A.)^65^. Further data analysis was performed using QIAGEN Ingenuity Pathway Analysis (QIAGEN Inc, Manchester, U.K., https://digitalinsights.qiagen.com/IPA)^66^. The raw data are deposited in the NCBI SRA (BioProject No: GSE288002).

### Statistical analysis

Statistical analysis was performed using one-way analysis of variance (ANOVA) with Dunnett’s multiple comparisons tests when comparing the means of 3 or more groups of data to the mean of the control group. Statistical analysis was performed using one-way ANOVA with Tukey’s multiple comparisons tests when comparing the means of 3 or more groups to the means of every other group. Statistical analysis was performed using paired t-tests when comparing the means of two groups of data. All statistical evaluations were performed using GraphPad Prism (Version 10.1.1) (San Diego, California, U.S.A.). Data is represented as the mean ± SD. Experimental n numbers and replicates are as described in figure legends. Significance was assessed in all experiments as a probability value of *P≤0.05, **P≤0.01, ***P≤0.001.

## Author contributions

Conceptualization, E.M.H., J.E.P.; Methodology, E.M.H., A.S.S., C.P. and R.P.A.; Formal Analysis, E.M.H.; Investigation, E.M.H.; Writing – Original Draft, E.M.H.; Writing – Reviewing & Editing, all authors; Visualization, E.M.H.; Funding Acquisition, R.K.B., J.E.P., M.P.P., and R.F.H.; Resources, A.S.S., R.P.A., M.P.P., and R.F.H.; Supervision, A.S.S, R.P.A., R.K.B., J.E.P., M.P.P., and R.F.H.

## Acknowledgements

EMH was funded by a Biotechnology and Biological Sciences Research Council (BBSRC) Case Studentship co-funded by Unilever (London Interdisciplinary Doctorate programme, LIDo), The authors would like to thank: the clinical team, Springfield Hospital Chelmsford, for providing redundant skin samples for skin cell isolation; Dr Sally Grimshaw, Unilever, for providing the microbial isolates; Dr Luke Gammon, QMUL Phenotypic Screening Facility, for his assistance with immunofluorescence imaging and analysis; and Dr Fei-Ling Lim, Unilever, for training and guidance on transcriptomic analysis.

## Data Availability

The data that support the findings of this study are available from the corresponding author, EMH, upon reasonable request.

## Supplementary Information

Document “McPolinHall_CellHostandMicrobe_supplementary”. Figures S1-S7 and Tables S1-S5.

## Resource Availability

### Lead contact

Requests for further information and resources should be directed to and will be fulfilled by the lead contact, Dr Esmé Poppy McPolin-Hall.

### Materials and availability

This study did not generate new unique reagents.

