## Supplementary for "Microbially-competent skin organotypics reveal dysregulated AhR signalling in microbial dysbiosis"

**Figure S1. Schematic depicting the generation and microbial inoculation of skin OTs using donor-matched hDFs, embedded in a collagen/matrigel matrix, and NHEKs.** OTs were cultured for 10 days prior to microbial colonisation and on day 11 were inoculated with microbes in PBS. Excess PBS was removed 3 hours after inoculation to maintain the air-liquid interface. OTs were cultured for 3 days post-microbial colonisation and media was changed daily. On day 14, OTs were harvested and cut in half, with half snap frozen and stored at -80°C for DNA/RNA extraction and half formalin fixed and paraffin embedded for H&E and IF analysis. Created using Biorender.

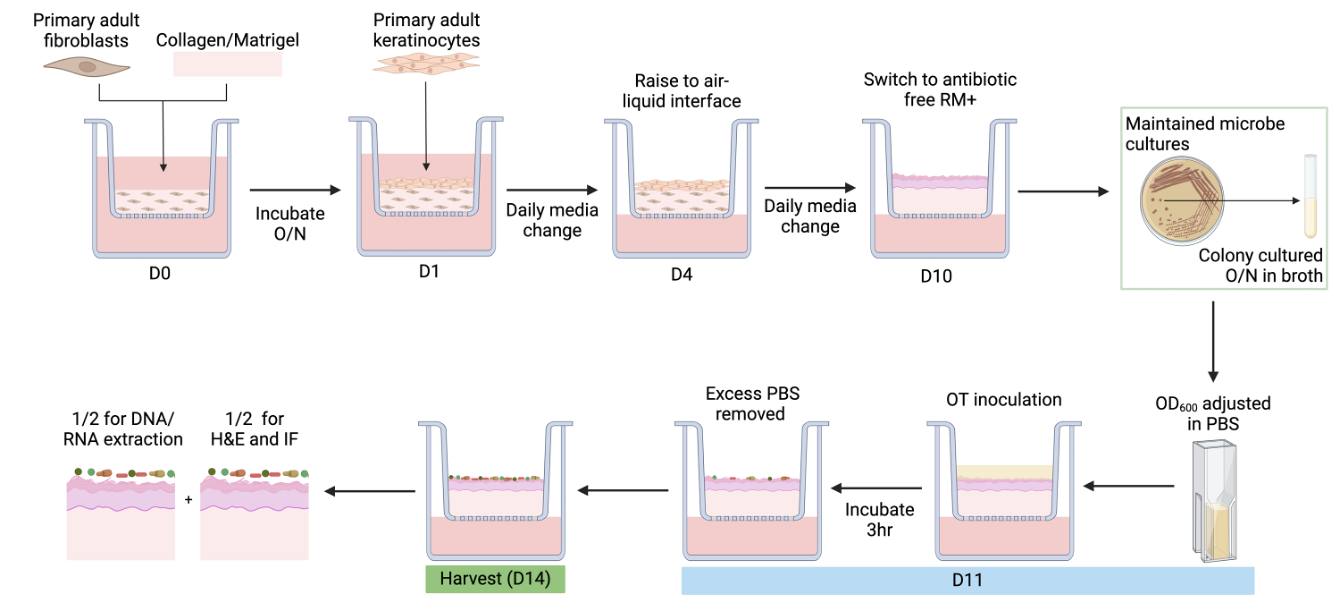

**Figure S2.** **Percentage cytotoxicity of NTERTs cocultured with serially diluted concentrations of microbes for 3 days, measured using Lactate Dehydrogenase (LDH) assay.** (A) To determine optimal seeding densities for inoculating OTs, serial dilutions of *S. epidermidis, S. capitis, C. acnes, M. restricta* and *M. globosa* were cocultured with NTERTs. The optimal seeding density of each microbial species was determined based on the seeding density that induced the lowest NTERT cytotoxicity whilst also enabling sufficient microbial biofilm formation. The red dotted line indicates the optimal seeding density determined. (B) Representative brightfield images of NTERTs cocultured with optimal seeding densities of each microbial species. Scale bar = 100μm. Data represents n=5 biological replicates, carried out in triplicate, ± SD.

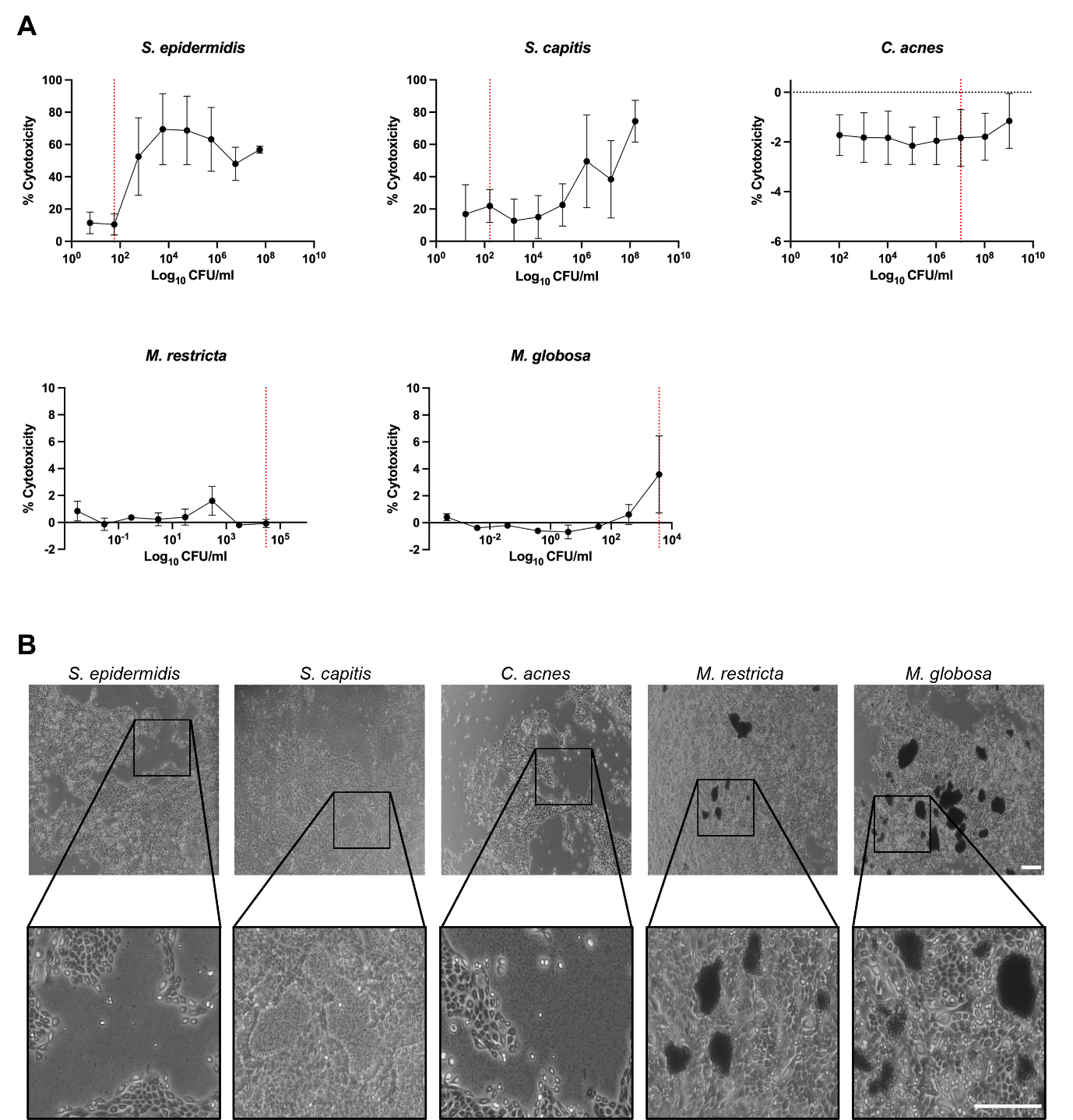

**Figure S3. H&E and IF images of human adult healthy scalp skin.** Redundant human adult healthy scalp skin was collected from redundant skin from elective rhytidectomy surgery. (A) Representative H&E image (Nanozoomer slide scanner, x20 magnification). Scale bar = 250µm. Representative IF images (INCA2200 widefield microscope) of human adult healthy scalp skin immunolabelled for (B) Keratin 10 (K10, green channel) (C) Keratin 14 (K14, red channel), (D) Ki67 (green channel) and (E) E-cadherin (Ecad, green channel). Cell nuclei were visualised using DAPI (blue channel). Data represents n=5 biological replicates. Scale bar = 100μm.

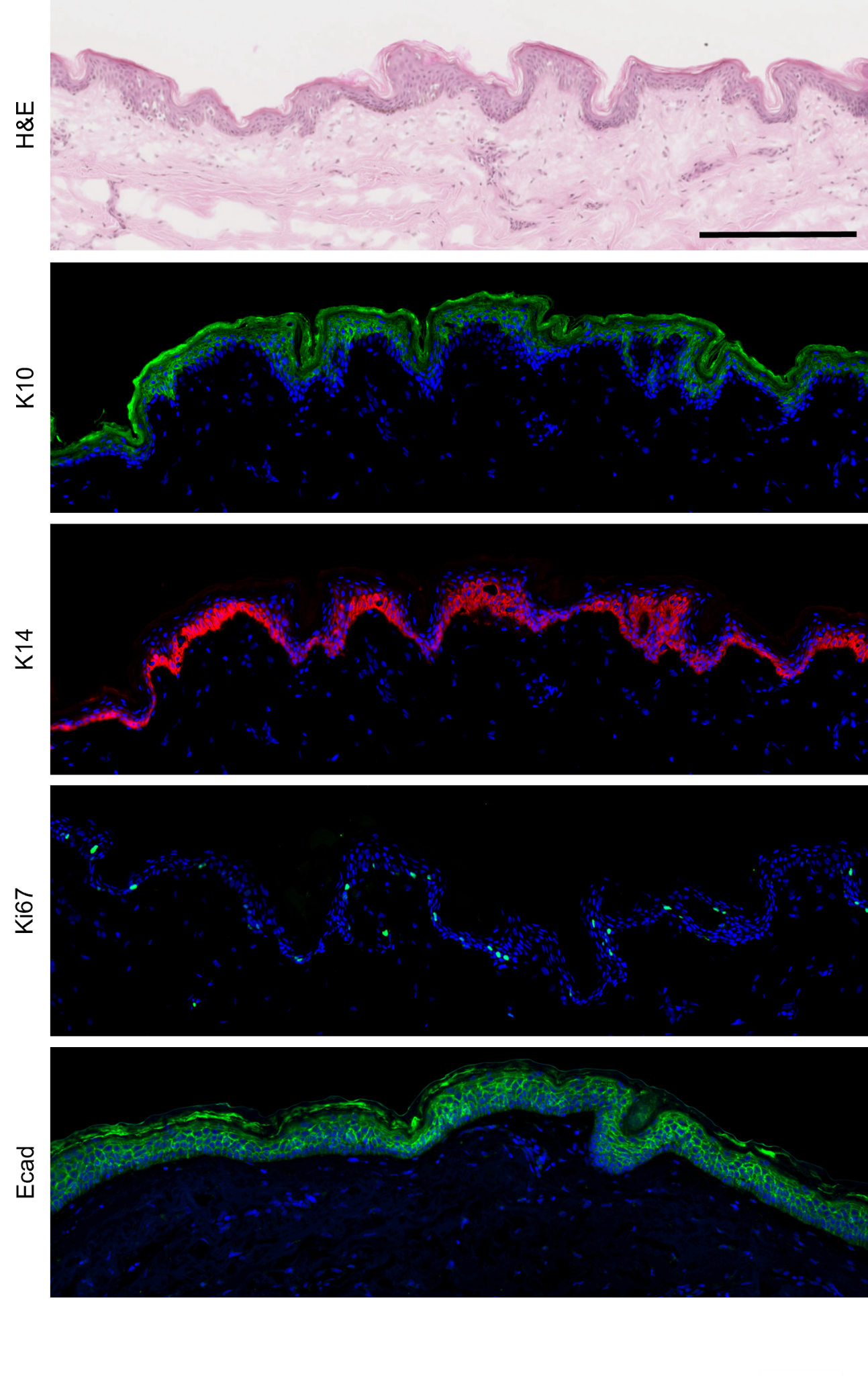

**Figure S4. IF analysis of sterile control, 5M colonised and 5MP colonised OTs.** Representative IF images (INCA2200 widefield microscope) of OTs immunolabelled for (A) K14 (red channel), (B) K10 (green channel), (C) Ki67 (red channel) and (D) E-cadherin (green channel) and quantification (HALO software) of FI and percentage of positive cells. Cell nuclei were visualised using DAPI (blue channel). Scale bar = 100µm. Data represents n=3 biological replicates carried out in duplicate ± SD. Sterile control (C), healthy microbiome inoculation (5M), dandruff microbiome inoculation (5MP). Statistical analysis was carried out using one-way ANOVA with Tukey’s multiple comparisons test. *p≤0.05, **p≤0.01, ***p≤0.001

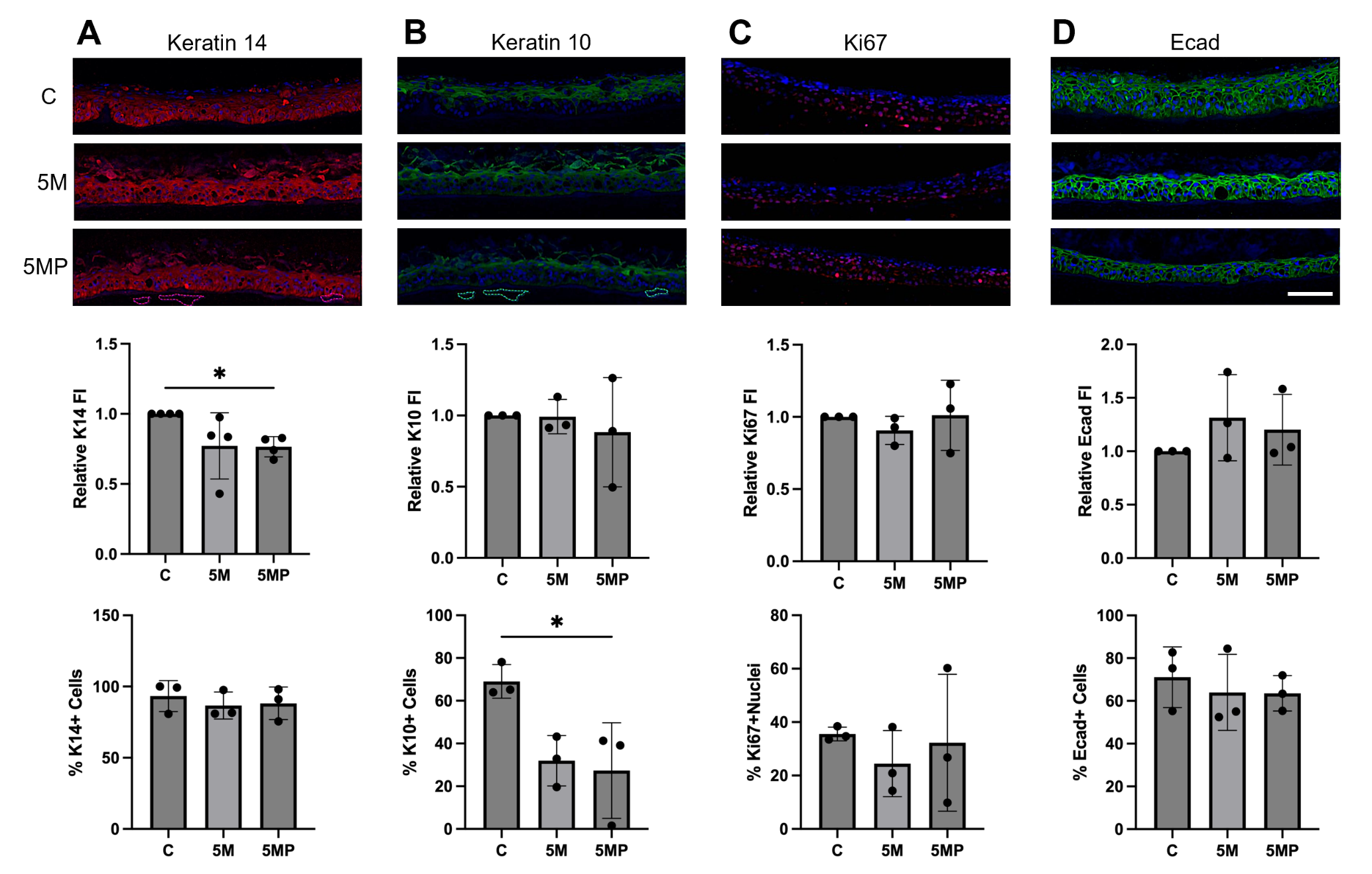

**Figure S5. The AhR signalling pathway.** Created using Biorender.

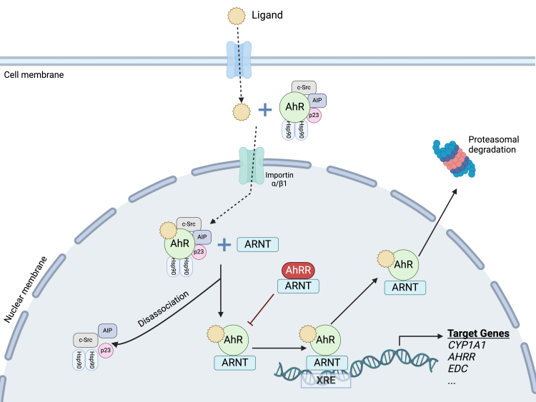

**Figure S6. OD_600_ vs CFU/ml standard calibration curves for *S. epidermidis, S. capitis, C. acnes, M. restricta* and *M. globosa.***

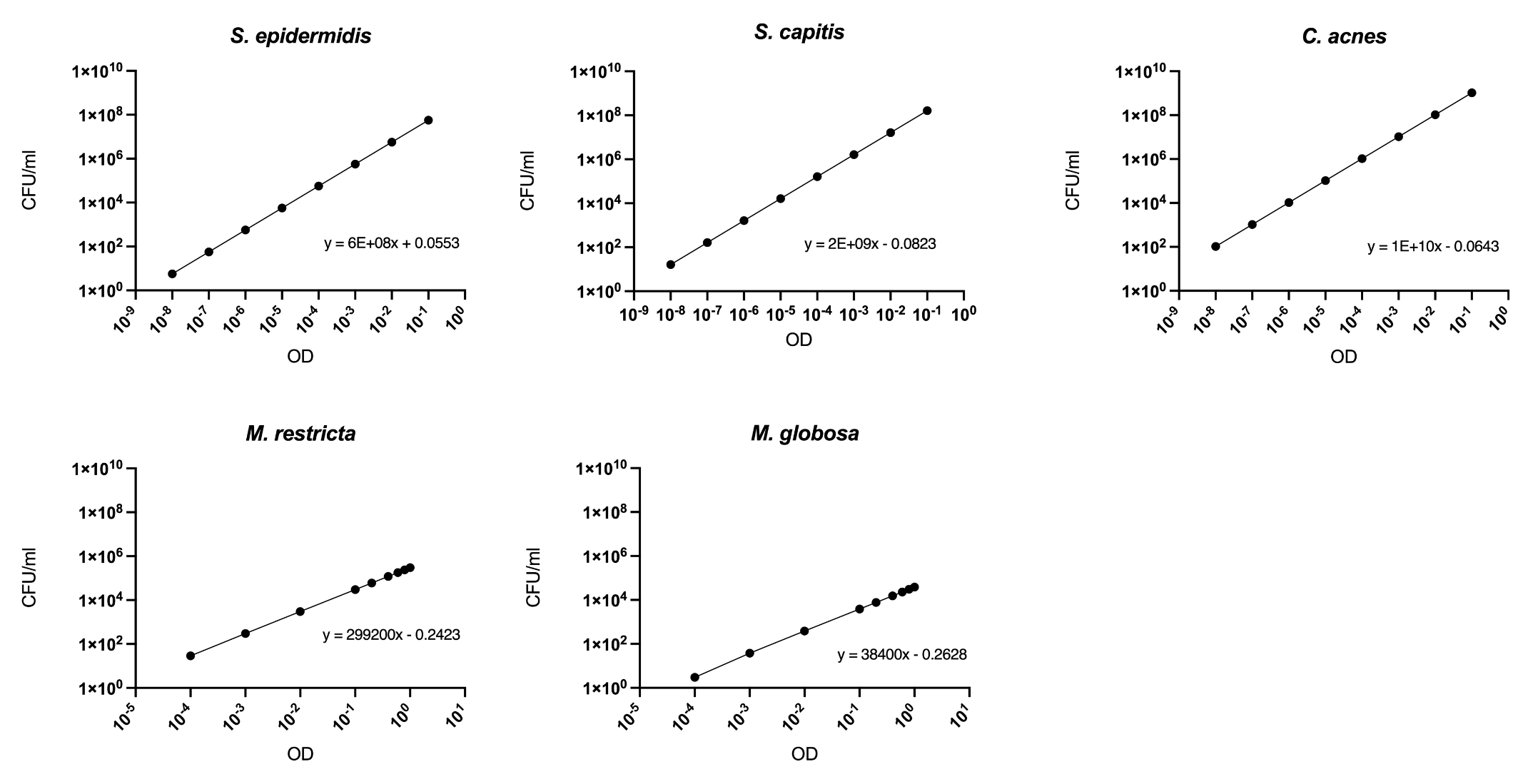

**Figure S7. RT-qPCR standard curves of Ct value vs Log_10_ DNA Copies/ml used to determine the quantity of genomic DNA of bacteria and fungi colonising OTs.**

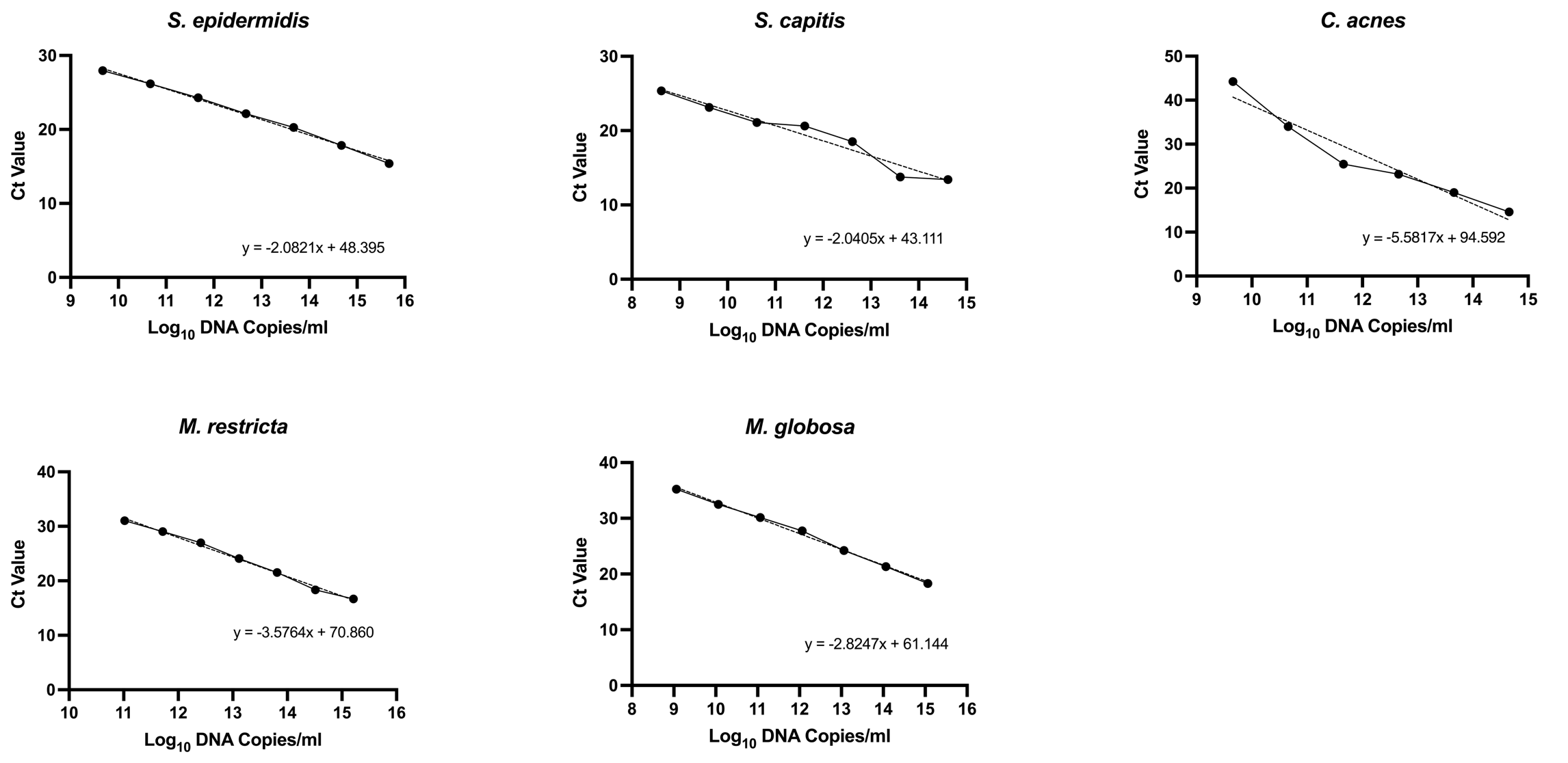

**Table S1. Table of species (and strain if applicable) used for microbiological culture, the agar and broth used for culture and culture conditions.**

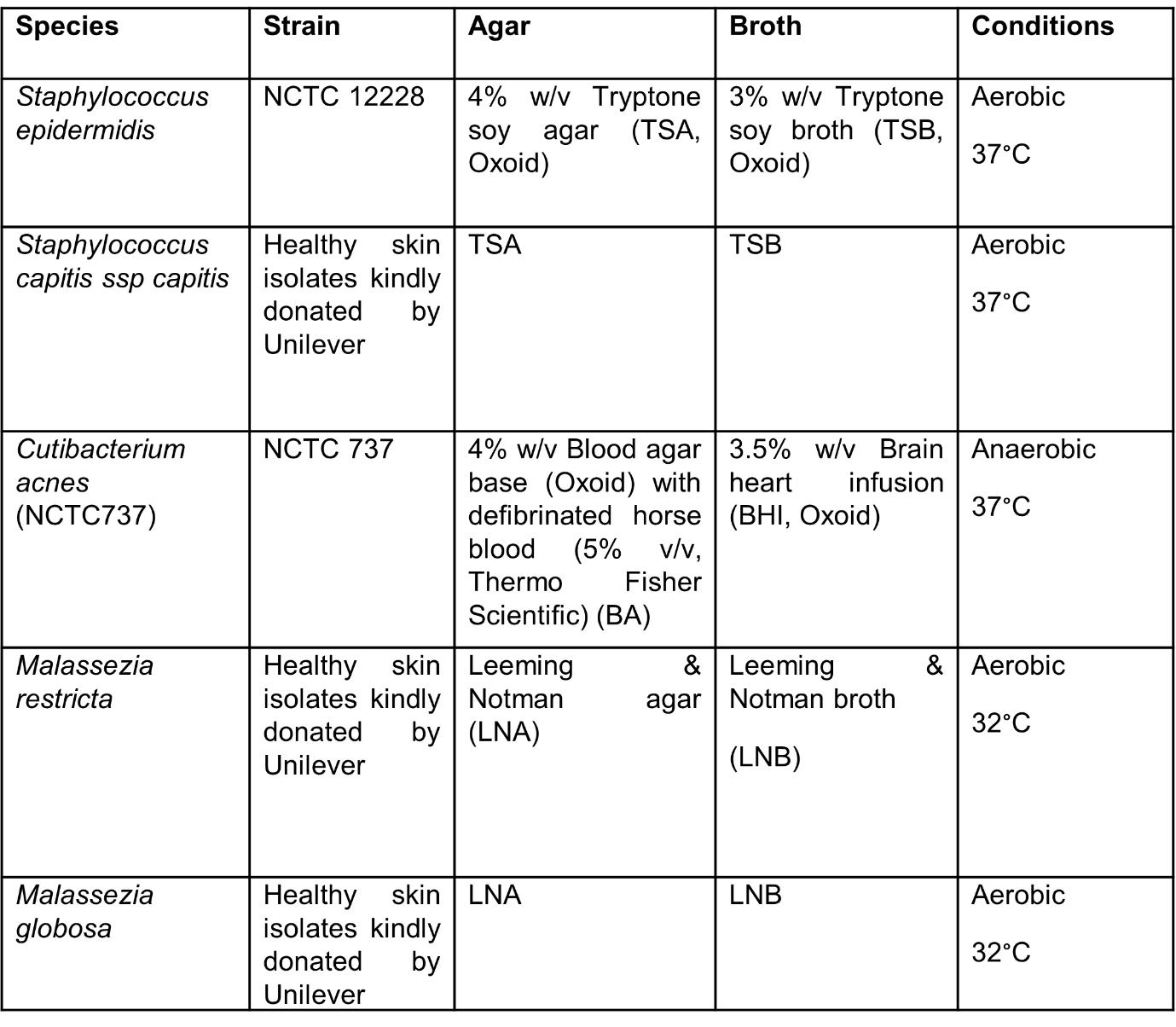

**Table S2. Primary and secondary antibodies used for immunofluorescence.** Antibody dilution in blocking buffer and subsequent final concentrations used. All primary antibodies are unconjugated, and all secondary antibodies are conjugated to Alexa Fluor™. The specific antigen retrieval buffer used for primary antibodies are indicated.

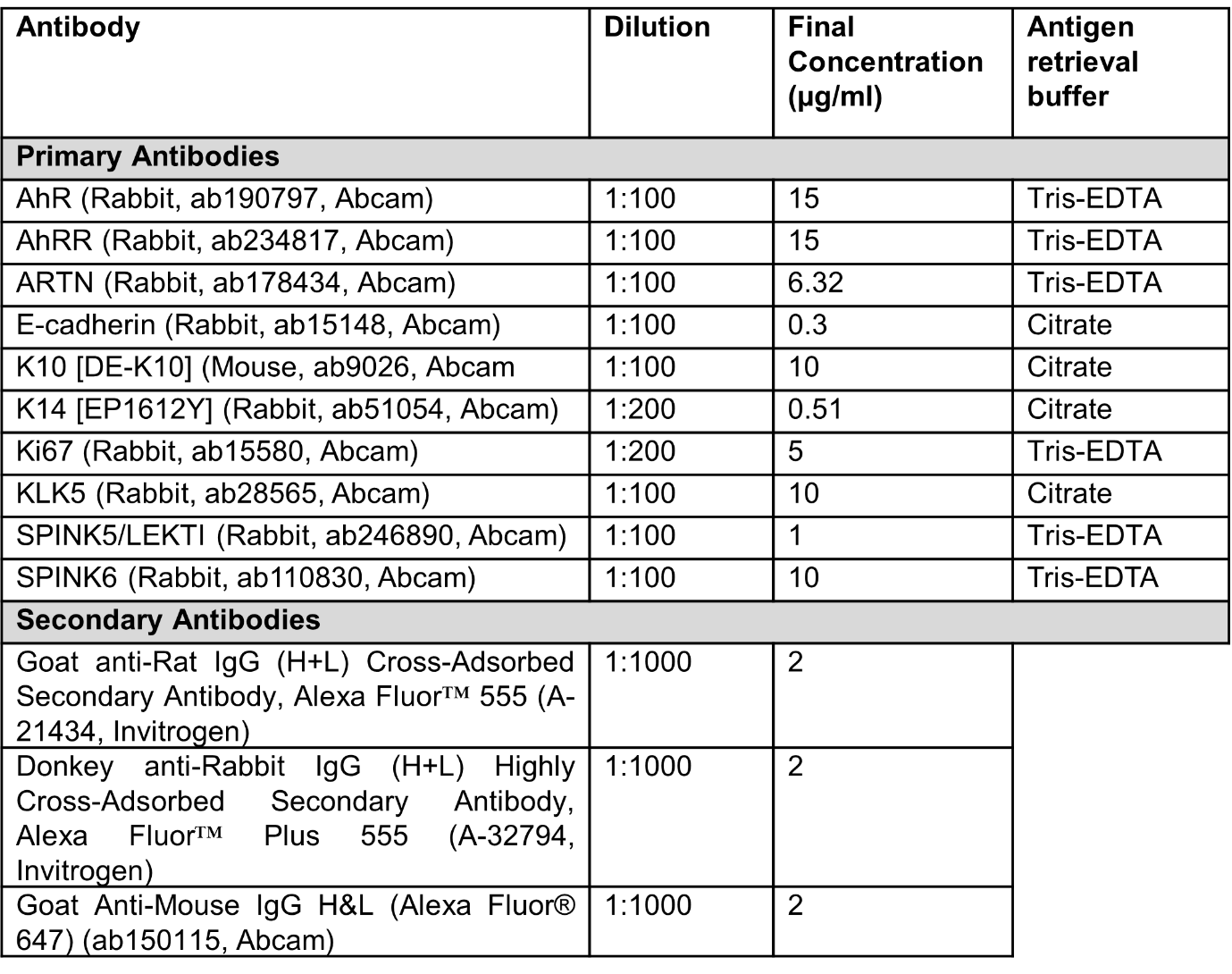

**Table S3.** **Sequences of qPCR primers for the identification and quantification of *S. epidermidis, S. capitis, C. acnes, M. restricta* and *M. globosa*.** FW = forward primer, RV = reverse primer.

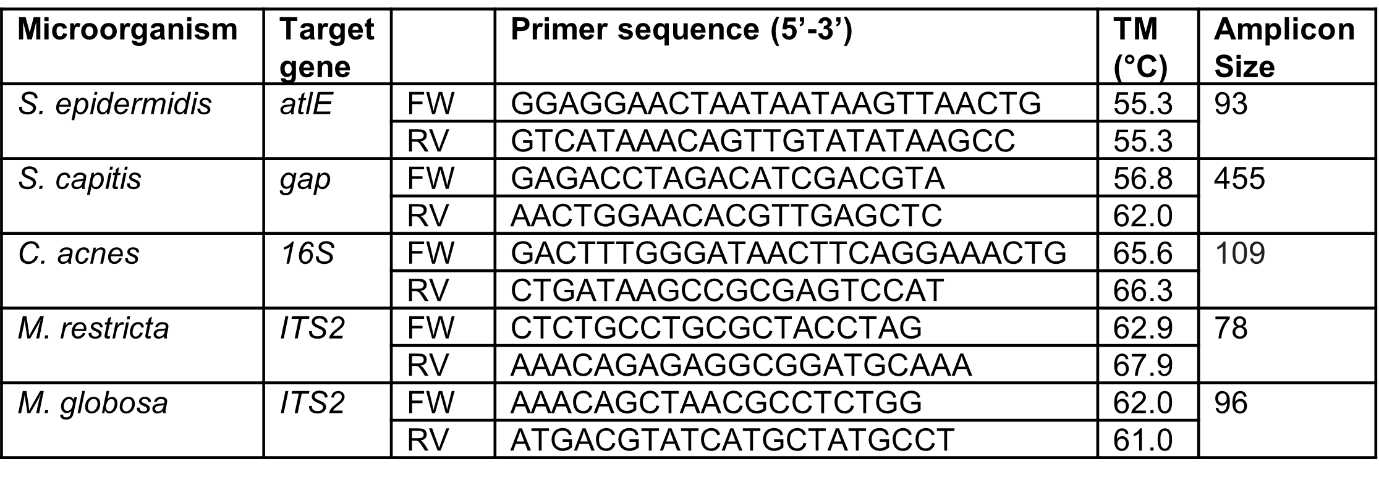

**Table S4. Sequences of qPCR probes for the identification and quantification of *S. epidermidis, S. capitis, C. acnes, M. restricta* and *M. globosa*.** Probes are dual-labelled with a 5’ fluorescent reporter dye (Fluorescein, 6-FAM) and a 3’ quencher (Black Hole Quencher-1, BHQ-1).

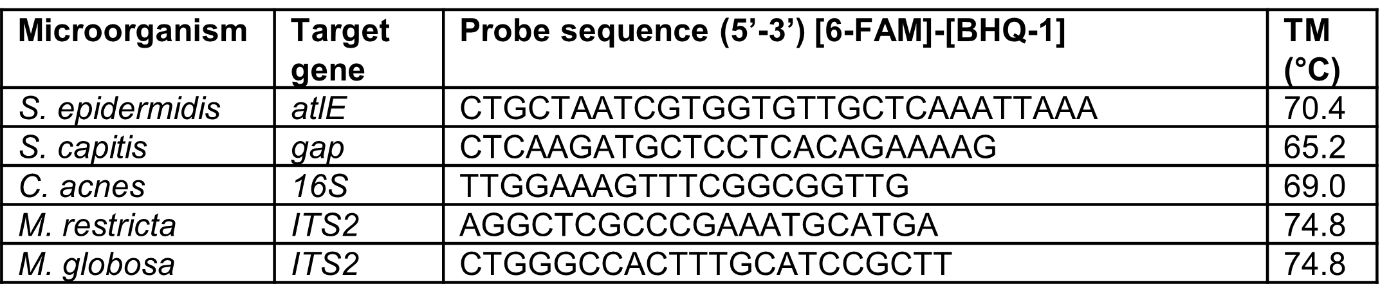

**Table S5. Gene Fragment sequences for each microbe, used to generate standard qPCR curves of known DNA copy number vs Ct value to quantify the abundance of microbes on OTs.** For fragment 1: red corresponds to *M. restricta* primer sites and yellow corresponds to *M. globosa* primer sites. For fragment 2: green corresponds to *S. epidermidis* primer sites and blue corresponds to *S. capitis* primer sites. For fragment 3: purple corresponds to *C. acnes* primer sites.

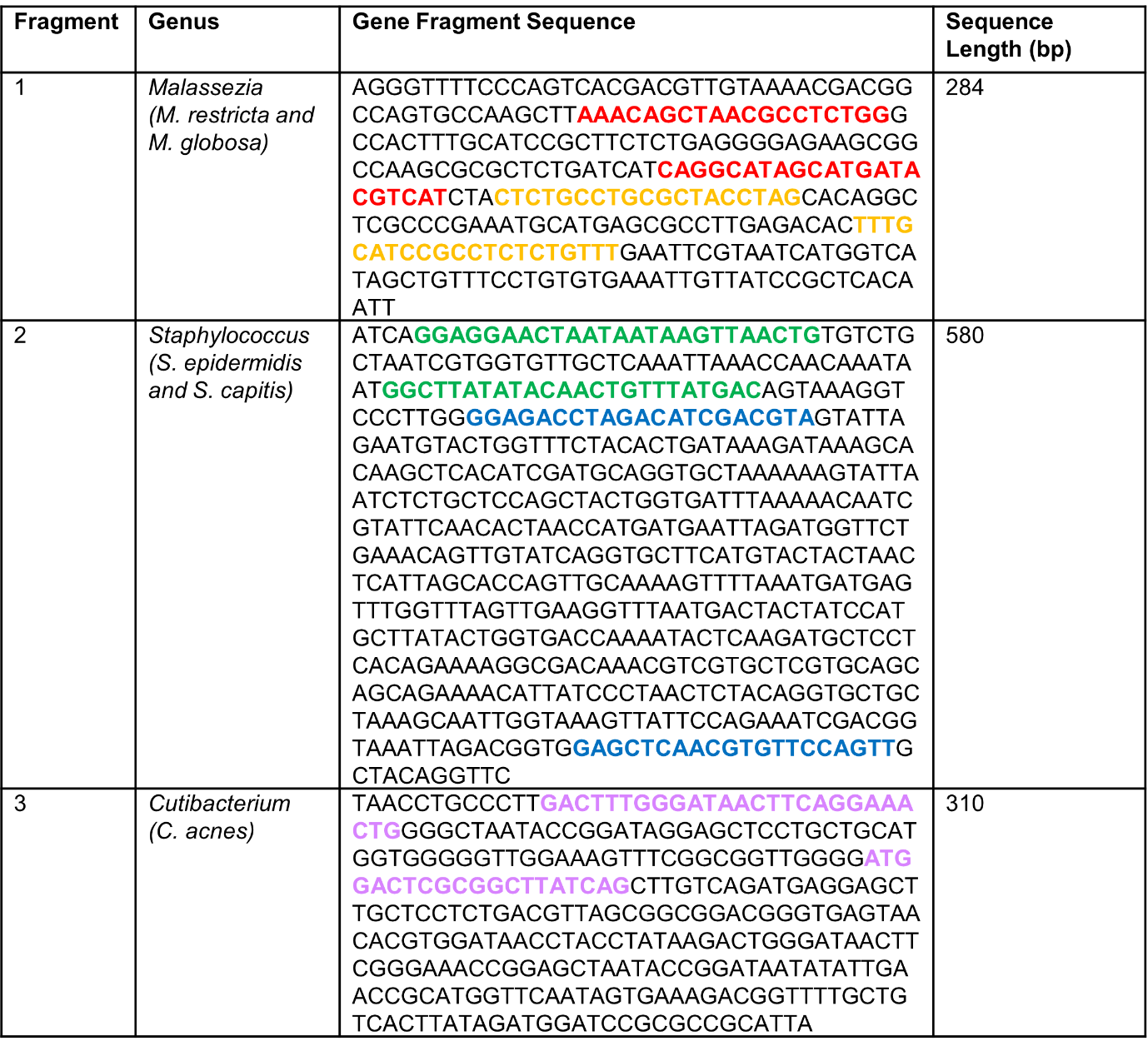
